# Enhanced Airway Epithelial Response to SARS-CoV-2 Infection in Children is Critically Tuned by the Cross-Talk Between Immune and Epithelial Cells

**DOI:** 10.1101/2023.05.17.541103

**Authors:** Vladimir G. Magalhães, Sören Lukassen, Maike Drechsler, Jennifer Loske, Sandy S. Burkart, Sandra Wüst, Eva-Maria Jacobsen, Jobst Röhmel, Marcus A. Mall, Klaus-Michael Debatin, Roland Eils, Stella Autenrieth, Aleš Janda, Irina Lehmann, Marco Binder

**Affiliations:** Research Group “Dynamics of Early Viral Infection and the Innate Antiviral Response”, Division Virus-Associated Carcinogenesis (F170), German Cancer Research Center (DKFZ), Heidelberg, Germany; Center for Digital Health, Berlin Institute of Health at the Charité - Universitätsmedizin Berlin; Faculty of Biosciences, Heidelberg University, Heidelberg, Germany; Molecular Epidemiology Unit, Center for Digital Health, Berlin Institute of Health at the Charité - Universitätsmedizin Berlin; Department of Pediatrics and Adolescent Medicine, Ulm University Medical Center, Ulm University, Ulm, Germany; Department of Pediatric Respiratory Medicine, Immunology and Critical Care Medicine, Charité – Universitätsmedizin Berlin, corporate member of Freie Universität Berlin and Humboldt-Universität zu Berlin, Berlin, Germany; German Center for Lung Research, associated partner, Berlin, Germany; Berlin Institute of Health at the Charité - Universitätsmedizin Berlin, Berlin, Germany; Health Data Science Unit, Faculty of Medicine, University of Heidelberg; Research Group “Dendritic Cells in Infection and Cancer” (F171), German Cancer Research Center (DKFZ), Heidelberg, Germany; University Hospital Tübingen, Department of Hematology, Oncology, Clinical Immunology and Rheumatology, Eberhard Karls University Tübingen, Tübingen, Germany

## Abstract

To cope with novel virus infections to which no prior adaptive immunity exists, the body strongly relies on the innate immune system. In such cases, including infections with SARS-CoV-2, children tend to fair better than adults. In the context of COVID-19, it became evident that a rapid interferon response at the site of primary infection is key for successful control of the virus and prevention of severe disease. The airway epithelium of children was shown to exhibit a primed state already at homeostasis and to respond particularly well to SARS-CoV-2 infection. However, the underlying mechanism for this priming remained elusive. Here we show that interactions between airway mucosal immune cells and epithelial cells are stronger in children, and via cytokine-mediated signaling lead to IRF-1-dependent upregulation of the viral sensors RIG-I and MDA5. Based on a cellular *in vitro* model we show that stimulated human peripheral blood mononuclear cells (PBMC) can induce a robust interferon-beta response towards SARS-CoV-2 in a lung epithelial cell line otherwise unresponsive to this virus. This is mediated by type I interferon, interferon-gamma and TNF, and requires induction of both, RIG-I and MDA5. In single cell-analysis of nasal swab samples the same cytokines are found to be elevated in mucosal immune cells of children, correlating with elevated epithelial expression of viral sensors. *In vitro* analysis of PBMC derived from healthy adolescents and adults confirm that immune cells of younger individuals show increased cytokine production and potential to prime epithelial cells. In co-culture with SARS-CoV-2-infected A549 cells, PBMC from adolescents significantly enhance the antiviral response. Taken together, our study suggests that higher numbers and a more vigorous activity of innate immune cells in the airway mucosa of children tune the set-point of the epithelial antiviral system. This likely is a major contributor to the robust immune response to SARS-CoV-2 in children. Our findings shed light on the molecular underpinnings of the stunning resilience of children towards severe COVID-19, and may propose a novel concept for immunoprophylactic treatments.

## Introduction

The ongoing COVID-19 pandemic represents one of the most devastating global health crises in modern times. COVID-19 is caused by the novel coronavirus SARS-CoV-2, with the first human infections detected in Wuhan, China, in late 2019. Despite most infected individuals develop only mild symptoms or remain asymptomatic, a notable fraction of patients develops a vast spectrum of severe disease, ranging from pneumonia and acute respiratory distress syndrome to multi-organ failure and death (1–3). World-wide, the virus has caused upwards of 6.5 million deaths (http://covid19.who.int, accessed Nov 4, 2022) with models suggesting a true death toll of up to 20 millions (4). While the determinants of severe disease are still not fully understood, chronic comorbidities such as hypertension, diabetes, cardiovascular disease and obesity as well as certain genetic factors clearly play a role (5). However, the clearest and most striking pre-disposing factor is age: the infection fatality rate of SARS-CoV-2 increases exponentially with age above 5 years (6). Compared to adults and in particularly elderly, children and adolescents generally display fewer symptoms with shorter duration, despite no strong difference in initial viral load (7, 8).

Hugely powered clinical association studies identified dysfunction of the antiviral type I interferon (IFN) system of the innate immune system to play a decisive role in the development of severe and life-threatening COVID-19 (9–11). This was seemingly at odds with numerous early reports describing exuberant production of cytokines (“cytokine storm”), including type I IFNs, as a hallmark of severe COVID-19 (3, 12, 13). However, these seemingly disparate findings can be reconciled when the kinetics of the IFN response is taken into account (14, 15): early and robust production of type I (and III) IFNs in the initially infected tissue of the upper respiratory tract is clearly antiviral and contributes to the immunological containment of the virus (11, 16, 17), whereas a delayed onset but lasting induction of IFN rather represents a secondary production of IFNs as a consequence of dysregulated immune responses (cytokine storm) and is associated with more severe courses of the disease (12, 13, 18). Recently, we and others have found a proper swift and local induction of IFN to be a key feature of the mucosal immune response to SARS-CoV-2 infection in children and adolescents, which likely is a major contributor to their striking resilience towards developing severe COVID-19 (19, 20).

SARS-CoV-2 is a positive-strand RNA virus that primarily infects ACE2-expressing ciliated epithelial cells in the lining of the respiratory tract (21), exhibiting rapid replication dynamics in permissive cells (18, 22, 23). Its ∼30 kb genome is among the largest known RNA virus genomes, encoding 29 proteins, many of which possess the capacity to antagonize antiviral immune responses (24, 25). Taken together, the immediate-onset translation (positive-stranded RNA), rapid RNA replication and the high number of immune antagonists render SARS-CoV-2 highly efficient in preventing the mounting of host cell-intrinsic antiviral responses, such as the production of type I and III IFNs (18, 23). This lack of IFN induction experimentally hampered the study of cellular sensing pathways, but recently the RIG-I-like receptor (RLR) MDA5 has been identified as the major sensor for SARS-CoV-2 in epithelial cells (26–28). Interestingly, downstream of MDA5, viral antagonists specifically block the activation of the transcription factor IRF3 and thereby the induction of IFNs, while the activation of NFκB remains possible, leading to the production of a pro-inflammatory set of cytokines (23); in fact, NFκB has been reported to support SARS-CoV-2 replication (29). Very likely, this skewed cytokine profile directly contributes to the dysregulation of downstream events such as the recruiting and activation of professional immune cells mediating a proper antiviral immune response (classically termed type 1 immunity) (15).

In the present study, we aimed at better understanding the underlying determinants for a proper induction of IFN in infected airway epithelial cells versus its failure. We previously observed a preactivated state of immune and epithelial cells in the upper airways of children prior to infection, characterized by a higher expression level of RLRs, in particular MDA5, as compared to adults (20). This sensitized state was associated with a rapid and robust induction of an IFN signature across epithelial and immune cell types upon SARS-CoV-2 infection. We hence speculate that subtle differences in the set-point of the virus sensing machinery in epithelial cells would have a decisive role in the successful mounting of an IFN response. Confirming this hypothesis, we report that priming of epithelial A549 cells with low to moderate doses of type I IFN or select inflammatory cytokines sensitizes their RLR pathway sufficiently to produce robust levels of type I and III IFNs upon SARS-CoV-2 infection. While MDA5 was the major sensor for this response, we additionally find a substantial contribution of RIG-I. With regard to why specifically children’s epithelial cells exhibit this primed state, we looked into the involvement of immune cells. In single cell RNA sequencing (scRNA-Seq) data of nasal swabs taken from healthy children or adults, we found a substantially higher number of immune cells in the nasal mucosa of children, as well as substantially stronger immune-epithelial interactions (20). In an *in vitro* model, we could show that microbially stimulated immune cells (PBMC) of children are able to induce stronger RLR expression in A549 epithelial cells than PBMC of adults, sufficient to enable robust IFN induction upon SARS-CoV-2 infection. This effect was mediated by both type I IFN as well as pro-inflammatory cytokines such as IFN-γ, IL-1β and TNF. We confirmed these cytokines are constitutively expressed by immune cells in the nasal mucosa of children, very likely conferring the priming of their epithelial cells and eventually enabling the robust immune response to SARS-CoV-2 infection observed in children and adolescents.

## Material and Methods

### Cell lines

A549, Calu-3, and HEK-293T cells were cultured in Dulbecco’s modified Eagle medium (DMEM high glucose, Life Technologies) supplemented with 10% (v/v) heat-inactivated fetal bovine serum (FBS) (Thermo Fisher Scientific), 100 µg/ml penicillin, 100 µg/ml streptomycin (Life Technologies), and 1% non-essential amino acids (Thermo Fischer Scientific), and maintained in a humidified atmosphere containing 5% CO2 at 37°C.

A549-derived knockout cell lines for MyD88 (30), MAVS (30), RIG-I (31), MDA5 (32), RIG-I/MDA5 (referred to as RIG-I/MDA5^DKO^)(32), IRF3 (33), IRF1 (34), IFNAR1(34), IFNAR1/IFNLR/IFNGR1 (referred to as IFNR^TKO^) (34) and non-targeting CRISPR control cells (NT) (34) were generated by CRISPR/Cas9 technology as described before. A549 knockout cell lines for TRIF and STING were generated using the lentiCRIPSR system (34, 35) with the following gRNA sequences: TRIF: 5’-GTGAGGCCAGGATCTCTCTAG-3’, STING: 5’-TGTGCGCAGCTCCTCAGCC-3’ were validated as shown in Suppl. Fig. 8.

A549 stably expressing RIG-I or MDA5 in RIG-I/MDA5^DKO^ cells were produced by lentiviral transduction using a pWPI-based vector, with transgene expression of codon-optimized RIG-I and MDA5 under the control of the murine ROSA26 promoter, which has a weak promoter activity in human cells. Empty vector controls were generated accordingly.

### Generation of PBMC

Peripheral blood mononuclear cells (PBMC) for bacterial stimulation were isolated from Buffy coats of healthy volunteers, obtained through ZKT Tübingen GmbH. Buffy coats were diluted 1:7 with Dulbecco’s PBS (Life Technologies). PBMC were obtained by density gradient centrifugation at 2000 rpm for 20 min at room temperature with 35 mL cell suspension stacked on 15 mL Biocoll separation solution (Biochrom). The interphase containing PBMC was abstracted and washed twice with PBS. PBMC were frozen in aliquots of 2×10^7^ cells in RPMI1640 containing 20% FBS and 10% DMSO at −150°C until further use. The study was approved by the local ethics committee (240/2018BO2) and complies with the declaration of Helsinki and the good clinical practice guidelines on the approximation of the laws, regulations and administrative provisions of the member states relating to the implementation of good clinical practice in the conduct of clinical trials on medicinal products for human use.

PBMC of donors of specific age were obtained from participants of the COVID-19 household study in Baden-Württemberg, Germany. This study was initiated by the University Children’s Hospitals in Freiburg, Heidelberg, Tübingen and Ulm and approved by the respective independent ethics committee of each center. Blood samples for this substudy were collected at study site Ulm in July 2020. Epidemiological and serological data describing the larger cohort have been published previously (36–38). The study is registered at the German Clinical Trials Register (DRKS), study ID 00021521, conducted according to the Declaration of Helsinki. For the present study, only donors were included that had no history of COVID-19 as well as had tested negative in three serological assays (EuroImmun-Anti-SARS-CoV-2 ELISA IgG /S1/, Siemens Healthineers SARS-CoV-2 IgG /RBD/, and Roche Elecsys Ig /Nucleocapsid Pan Ig/). PBMC were selected from heparinized full blood by a standard density gradient (Pancoll Separating Solution, PAN-Biotech), and cryopreserved in medium containing 75% FBS, 15% RPMI (PAN-Biotech) and 10% DMSO (Serva) at −196°C in liquid nitrogen until further analysis.

### Cytokine stimulation

A549 IFNAR1^KO^, IFNR^TKO^, and the corresponding NT control cells were seeded at a density of 1.5 x 10^5^ cells per well in 500 µl supplemented DMEM. The next day, cells were stimulated with 10 ng/ml of TNF (ab259410, Abcam), IL-6 (7270-IL, R&D Systems), IL-1β (ALX-502-001, Enzo Life Sciences), IL-10 (ab259402, Abcam) and IFN-γ (285-IF-100/CF, R&D Systems). After 8 hours, cells were washed with PBS and lysed by adding 300 µl Monarch RNA Lysis Buffer and stored frozen −80°C until further analysis.

### Infection

For infection with SARS-CoV-2, A549 cells were seeded in a 24-well plate at a density of 4×10^4^ cells. The next day, cells were transiently transduced with lentiviral vectors coding for ACE2 and TMPRSS2. In order to ensure homogeneous high efficiency transduction, A549 cell were incubated with lentiviral supernatants diluted 1:20 (v/v) in culture medium containing 10 µg/ml polybrene (Merck Millipore) and centrifuged at 805xg at 20 °C for 30 min. At 24 hours post-transduction, cells were either mock-stimulated or simulated with 170 U/ml IFN-β or 10 ng/ml TNF for 16 and 8 hours, respectively. After stimulus, cells were washed once with PBS and incubated with 200 µl/well of DMEM containing SARS-CoV-2 at an MOI of 0.1 (IFN-β) and 1 (TNF). One hour after inoculation, cells were washed once with PBS and fed with 500 µl/well of DMEM supplement with 2% FCS. 24 hours after inoculation, cells were washed once with PBS and lysed for qPCR and western blot analyses.

### RNA isolation and RT-qPCR

Total RNA was isolated from cells using the Monarch Total RNA Miniprep Kit (New England Biolabs), employing on-column digestion of residual DNA by DNAseI treatment, according to the manufacturer’s instructions. Isolated RNA was reverse transcribed using the High Capacity cDNA Reverse Transcription Kit (Thermo Fisher Scientific) with random hexamer primers according to the manufacturer’s specifications. Transcript levels were assessed by real-time PCR using the iTAQ Universal SYBR Green Supermix (BioRad) on a BioRad CFX 96 Real-Time PCR Detection System. Primers for qPCR were used as the following: SARS-CoV-2-ORF1 forward: 5’-GAGAGCCTTGTCCCTGGTTT-3’, SARS-CoV-2-ORF1 reverse: 5’-AGTCTCCAAAGCCACGTACG-3’, IFIH1(MDA5) forward: 5’-TCGTCAAACAGGAAACAATGA-3’, IFIH1 reverse: 5’-GTTATTCTCCATGCCCCAGA-3’, RIGI (RIG-I) forward: 5’-CCCTGGTTTAGGGAGGAAGA-3’, RIGI reverse: 5’-TCCCAACTTTCAATGGCTTC-3’, IFNB1 (IFN-β) forward: 5’-CGCCGCATTGACCATCTA-3’, IFNB1 reverse: 5’-GACATTAGCCAGGAGGTTCTC-3’, GAPDH forward: 5’-CGGAGTCAACGGATTTGGT-3’, GAPDH reverse: 5’-TTCCCGTTCTCAGCCTTGAC-3’, 45S rRNA forward: 5’-GAACGGTGGTGTGTCGTT-3’, 45S rRNA reverse: 5’ -GCGTCTCGTCTCGTCTCACT-3’, RSAD2 forward: 5’-CGTGAGCATCGTGAGCAATG-3’, RSAD2 reverse: 5’-CTTCTTTCCTTGGCCACGG-3’, IFIT1 forward: 5’-GAATAGCCAGATCTCAGAGGAGC-3’, IFIT1 reverse: 5’-CCATTTGTACTCATGGTTGCTGT-3’. Quantification of target gene expression was done using the ΔCT method (normalization to housekeeping gene only) or ΔΔCT method (normalization to housekeeping gene and to a reference cell line / condition).

### Western blotting

Mock and infected samples were lysed with 100 µl sample buffer (30 mM Tris [pH 6.8], 0.05% bromophenol blue, 10% glycerol, 1% SDS, 2.5% β-mercaptoethanol) supplemented with 1 µl benzonase to cleave contaminant DNA. Denaturation was obtained by heat at 95°C for 5 min. Samples were subjected to SDS-PAGE on a 10% polyacrylamide gel for 90 min at 100V and blotted onto a polyvinylidene (PVDF) membrane (0.2 µm) by wet transfer for 120 min at 350 mA. Membranes were blocked with blocking buffer (5% BSA in TBS containing 0.1% Tween 20) for 1 hour at room temperature, followed by overnight incubation with primary antibodies on a shaker at 4°C. After washes with TBS containing 0.1% Tween (TBS-T), membranes were incubated with horse radish peroxidase (HRP)-conjugated secondary antibody for 1 hour in blocking buffer at room temperature. Membranes were rewashed 3x with TBS-T, and proteins were visualized using the Clarity Western ECL substrate (Bio-Rad) and a high-sensitive charge-coupled-device (CCD) camera (ChemoCam Imager 3.2; INTAS). Images were processed by the Imagej/Fiji software. The following commercially available antibodies were used: rabbit monoclonal antibodies anti-p-IRF3(S396) (#4947S), anti p-STAT1 (Y701) (#7649S), anti-STAT2 (#72604S), anti p-STAT2(Y690) (#D3P2P) were purchased from Cell Signaling Technology. Mouse monoclonal antibodies anti-GAPDH (sc-47724) and anti-IRF3 (sc-3361) were from Santa Cruz Biotechnology. Mouse monoclonal anti-RIG-I (AG-20B-0009) and anti-STAT1 (610115) were obtained from Adipogen and BD Biosciences, respectively. Rabbit polyclonal anti-MDA5 (ALX-210-935) was purchased from Enzo Life Sciences and mouse monoclonal anti-β-actin (A5441), along with anti-mouse and anti-rabbit antibodies coupled with HRP (A4416, A6154) were obtained from Sigma-Aldrich.

### Stimulation and infection of PBMC

Isolated PBMC were thawed in a 37°C water bath, mixed with 9 ml culture medium supplemented with 50 KU DNAse I (Millipore), and centrifuged at 350xg for 10 min. The supernatant was then aspirated and cells resuspended in 5 ml culture medium containing 200 KU DNAseI, kept at 4 °C for 20 min and viable cells quantified by trypan blue. PBMC were seeded at 1×10^6^/ml into 24-well tissue plates. For stimulation with *Yersinia enterocolitica* (WA-314, serotype O:8), the bacterial stock was pelleted, washed once with PBS, resuspended in culture medium and mixed with PMBC at a ratio of one bacterium per cell for 1 h. Bacterial growth was inhibited by supplementing media with 20 µg/ml gentamicin (Biochrom). At 24 hours post-treatment, cell suspensions were collected, centrifuged at 1,000xg for 5 min and the resultant supernatants were frozen at −80°C until further analysis.

### Co-culture of A549 and PBMC

For co-culture experiments, A549 ^ACE2/TMPRSS2^ were mock infected or infected with SARS-CoV-2 for 8 hours, followed by incubation with 1×10^6^ isolated PBMC in 1 ml culture medium in a 24-well plate. Cell suspensions were collected after 16 hours co-culture, centrifuged at 1,000xg for 5 min and supernatants were harvested and treated with 0.1% beta-propiolactone at room temperature for 1 hour and at 37°C for 2 hours. Samples were stored at −80°C until further analysis.

### dsRNA stimulation via electroporation

To synchronously stimulate the RIG-I pathway in A549 cells, electro-transfection was used as described in (39). Briefly, 4×10^6^ mock-treated or IFN-treated cells were resuspended in 400 µl cytomix buffer (120 mM KCl, 0.15 mM CaCl2, 10 mM KPO4, 25 mM HEPES, 2 mM EGTA, 2 mM MgCl2) and transferred to a 0.4 cm cuvette chamber containing 220 ng of *in vitro* generated 400-bp 5’ ppp-dsRNA (40). The suspension was electroporated using the Gene Pulser Xcell modular system (150 V, 10 ms exponential decay). Electroporated cells were transferred to pre-warmed DMEM and washed twice with DMEM before distributing them on 6-well plates containing 5×10^5^ cells in 1.2 ml medium per well. At the indicated time points, cells were washed once with PBS and lysed for qPCR analysis.

### Read-out by MSD Electrochemiluminescent Multiplex Assay

To quantify the level of released IFN, chemokine and proinflammatory cytokines, supernatants of mock and stimulated PBMCs were subjected to electrochemiluminescent multiplex assay using kits from Meso Scale Discovery. IFN-α2, IFN-β, IFN-λ1 (IL-29), and IFN-γ were measured with human U-PLEX Interferon Combo (K15094K), whereas the proinflammatory cytokines IL1-β, IL-6 and TNF were determined using the human V-PLEX Proinflammatory Panel 1 Kit (K15049D). Production of GM-CSF, IL-2, IL-4, IL-8, IL-10, and IL12p70 were detected by a customized U-PLEX Biomarker Group 1 (K15067L). Assays were performed according to the manufacturer’s instructions, read out by the MSD Quickplex SQ120 device and data evaluated using the MSD Discovery Workbench software (version 4.0). Values below the limits of quantification (LLoQ) were denoted as not detectable (nd).

### Single Cell RNA Sequencing Data Analysis

Single cell data including details on sample preparation, study participants (exact ages) and on the molecular definition of cell types and states was previously previously (20) and is publicly available as a Seurat object (doi: 10.6084/m9.figshare.14938755). Briefly, nasal airway swabs of healthy and of SARS-SoV-2-infected children and adults were obtained and analyzed by single cell RNA sequencing (scRNA-Seq). For the present study, further analyses were performed, exclusively based on the samples of healthy donors, comprising 18 children (4.03 to 16.11 years, median age 9.0 years) and 23 adults (24 to 77 years, median 46.0 years). Therefore, the Seurat object was subsetted to only SARS-CoV-2 negative samples using the subset function of Seurat (https://github.com/satijalab/seurat). The final object had 114,296 cells (SARS-CoV-2 negative children 51,595; negative adults 62,701). For all scRNA-Seq analyses Seurat 3.2.2 was used.

Cell–cell interaction analyses of scRNA-Seq data were performed using CellChat 1.5.0 (41). The interaction database was augmented with novel interaction pairs identified by Shilts et al. (42). Only interactions emanating from immune cells, i.e. with ligands being expressed by immune cells, and targeting receptor-expressing epithelial cells were retained. Pathway-level interactions were derived using the built-in annotations of CellChat.

### Statistical analysis

Unless stated otherwise, experiments have been performed three times independently, with each repetition comprising technical duplicates or triplicates. Shown are mean values with error bars representing standard deviations (SD), always representing the means and SD across independent repetitions (with technical replicates averaged before). For the sake of transparency, means of each repetition are displayed as individual data points in the graphs.

Differences between two data sets were tested for statistical significance by unpaired, one-tailed Student’s t-test if not stated otherwise. We used one-tailed tests as all experiments study effects with a pre-defined direction (*induction* of gene expression/cytokine production or *inhibition* of virus replication) and the null hypothesis was formulated accordingly. Statistical analyses and data plotting were performed using GraphPad Prism (v.9).

P-values for violin plots showing MDA5 and RIG-I expression levels across all epithelial cells were calculated using the Seurat function FindMarkers() applying the Wilcoxon test and adjusted with Bonferroni correction. Age-groups were chosen such that they reflect the actual ages of participants (4-7: n= 5, 8-11: n=8, 12-16: n=5); binning in the classical 0-6, 7-10, 11-18 groups did not change statistical significance.

P-values for violin plots showing average cytokine expression in immune cells across samples were calculated using the ks.test function of R and corrected with the Benjamini-Hochberg method (43).

## Results

### Priming with type I IFN renders lung epithelial cells competent to respond to SARS-CoV-2 infection

SARS-CoV-2 infection has been described to elicit a severely skewed cytokine response in epithelial cells, with a substantially dampened IFN component but high amounts of pro-inflammatory mediators(18, 21, 23, 44) (for a scheme of the cell-intrinsic antiviral pathways relevant for this study, see Fig. 1A). This lack of an early antiviral innate immune response at the site of primary infection is thought to be a major contributor to the development of severe COVID-19 (19). We and others have observed swifter and stronger IFN-responses upon SARS-CoV-2 infection in the upper airways of children than in those of adults (19, 20, 45, 46). In scRNA-Seq analyses, we found this to correlate with the presence of a subtle IFN-like transcriptional signature in nasal epithelial cells of children even in the absence of infection, with a clearly higher expression of genes of the viral sensing machinery such as RIG-I (encoded by the *RIGI* gene) and MDA5 (encoded by the *IFIH1* gene) (20). This suggested a “primed” state of epithelial cells in children rendering their innate antiviral system more sensitive in detection and more robust in its response.

**Figure 1.**
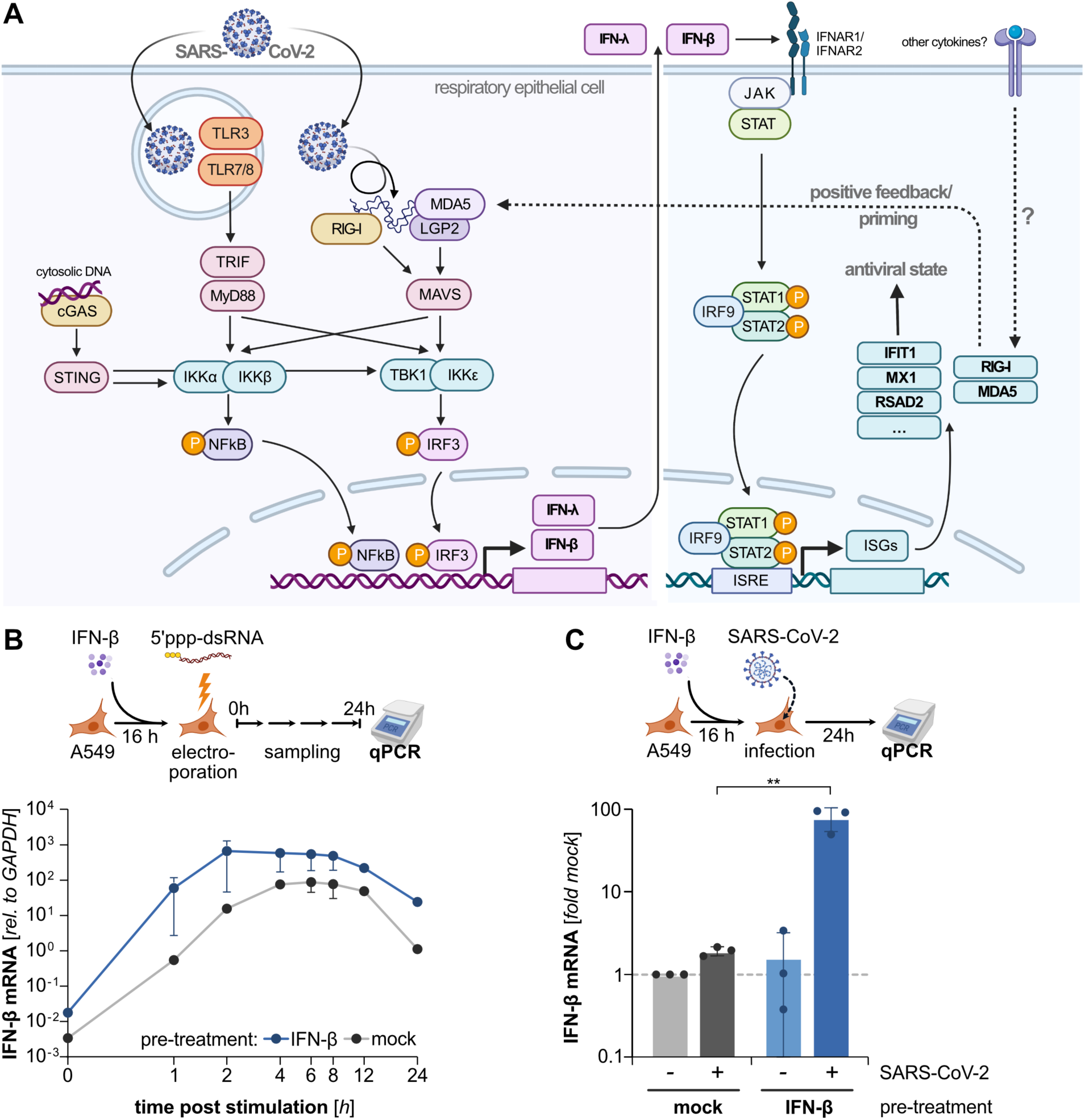
Impact of cell priming on RLR signaling kinetics and interferon response to SARS-CoV-2 infection. **(A)** Schematic overview of the cell-intrinsic antiviral response pathways relevant to this study. Pattern recognition receptors detect non-self nucleic acids triggering the transcription and production of type I and III IFN (left). Secreted IFN stimulates JAK/STAT signaling in an autocrine and paracrine manner. IFN stimulated genes (ISGs) include pattern recognition receptors RIG-I and MDA5 (right). **(B)** A549 cells were mock-treated or primed with 1xEC50 (170 IU/ml) of IFN-β for 16 hours. RIG-I was then synchronously stimulated in all cells by electro-transfection of 5’ppp-dsRNA. At the indicated time points, cells were lysed and analyzed for IFN-β (*IFNB1*) gene expression by qRT-PCR. Shown are means ±SEM of two biologically independent experiments. **(C)** A549^ACE2/TMPRSS2^ cells were mock-treated or primed as in (A), followed by infection with SARS-CoV-2 (MOI 0.1). At 24 hours post infection, IFN-β expression was measured by qRT-PCR. Values were normalized to 45S rRNA levels and expressed relative to non-infected mock-treated cells. Bars represent means ±SD of three independent experiments (individual experiments shown as dots). Statistical significance was tested by an unpaired two-tailed t-test; ** p<0.01.

To determine the cellular and molecular underpinnings of the “innate priming” of epithelial cells, we simulated the primed state and assessed its effect on antiviral signaling dynamics by using a moderate dose of type I IFN to pre-stimulate A549 human lung epithelial cells. We employed 170 IU/ml of IFN-β, determined to be the EC_50_ with regard to ISG induction in A549 cells (Suppl Fig 1A), and pre-treated cells overnight (16 hours). We then triggered RLR-signaling by electro-transfection of the RIG-I agonist 5’ppp-dsRNA, as described in (39). To monitor RLR pathway activity over time, we measured IFN-β (encoded by the *IFNB1* gene) transcription as a specific downstream readout for antiviral PRR signaling. In non-pre-treated cells, dsRNA stimulation gradually induced IFN-β expression, reaching maximum levels at 4-8 hours and declining thereafter (Figure 1B, gray line). As expected, pre-treatment alone did not trigger notable IFN-β transcription (see also Fig. 1C). However, upon dsRNA stimulation, IFN-β primed cells mounted a 1-2 log stronger response than non-pre-treated cells (Fig. 1B, blue line). This response was not only higher in amplitude but also notably swifter, reaching peak activation already at 2 hours after stimulation.

While priming of cells with IFN is not a novel concept, we propose that in particular this kinetic impact– often neglected in previous studies – will provide a significant advantage for cells in responding to and fending off virus infection. Many viruses, and in particular also SARS-CoV-2, encode and express antagonists of the cellular antiviral defense (25). We and others have described SARS-CoV-2 to potently suppress the induction of IFNs in respiratory epithelial cells likely due to its rapid replication and expression of a multitude of antagonists (23, 25, 47). We hypothesize that upon priming, the faster response dynamics of the antiviral system could confer a substantial advantage (“head-start”) to cells, such that the IFN system could be elicited before viral antagonism is fully mounted. To test this hypothesis, we pre-treated A549 cells expressing ACE2 and TMPRSS2 (A549^ACE2/TMPRSS2^) with the same regimen of 1x EC50 of IFN-β for 16 hours and then infected them with SARS-CoV-2. As described previously, SARS-CoV-2 infection did not elicit an IFN-β response in mock-treated A549 cells (Fig. 1C, gray bars). In contrast, IFN-priming of cells restored their capacity to mount a robust response upon SARS-CoV-2 infection with ∼100-fold induction of IFNB1 transcripts (Fig. 1C, blue bars). We confirmed these observations also in another human lung epithelial cell line, Calu-3, in which IFN levels ensuing SARS-CoV-2 infection were also increased upon prior priming (Suppl. Fig. 1B). Taken together, our data highlight the importance of the *relative* dynamics of viral replication *versus* host responses in determining the functional outcome of infection. This demonstrates that tissue exposure to cytokines can critically modulate the responsiveness of epithelial cells against SARS-CoV-2.

### Primed epithelial cells mount an effective antiviral state upon SARS-CoV-2 infection through viral sensing by MDA5 and RIG-I

Next, we wondered which PRR(s) detect SARS-CoV-2 in primed epithelial cells. Previous studies pinpointed MDA5 as its major cytosolic sensor (26–28, 48), but also TLRs have been implicated in cytokine responses to the virus (49, 50). We, therefore, made use of A549 cells with CRISPR/Cas9-based functional knockouts (KO) of the central adaptors of different PRR axes: MyD88 and TRIF for the TLRs, STING for cytosolic DNA sensors such as cGAS, and MAVS for the RLRs RIG-I and MDA5 (see Fig. 1A for a schematic overview). We observed that only cells devoid of MAVS entirely abrogated IFN-β induction, while KO of STING, MyD88, or TRIF had no effect (Figure 2A). To identify the actually responsible receptor upstream of MAVS, we investigated the role of RIG-I and MDA5 individually by priming individual KOs and a double KO cell lines and assessing their IFN response upon infection. Since MDA5 and RIG-I are classical ISGs themselves, pre-treatment of cells with IFN-β resulted in high MDA5 and RIG-I protein levels (Suppl. Fig. 2A; also Fig. 2C, compare NT and IFNAR^KO^). SARS-CoV-2 infection of primed cells lacking RIG-I but expressing high levels of MDA5 exhibited strong IFN-β induction similar to or even slightly stronger than wild-type (NT) cells (Fig. 2B). In contrast, absence of MDA5, despite high levels of RIG-I, resulted in a 2.4-fold reduction of IFNB1 transcripts relative to control cells. Strikingly, IFN-β induction was fully abolished when both sensors were lacking (Fig. 2B), suggesting that both RIG-I and MDA5 are essential for a full-fledged IFN response in epithelial cells. This was also confirmed at the downstream level of STAT2 activation, at which phosphorylation was still observed upon infection of RIG-I or MDA5 single KOs with SARS-CoV-2, but was fully diminished only in the double KO cells (Fig 2C).

**Figure 2.**
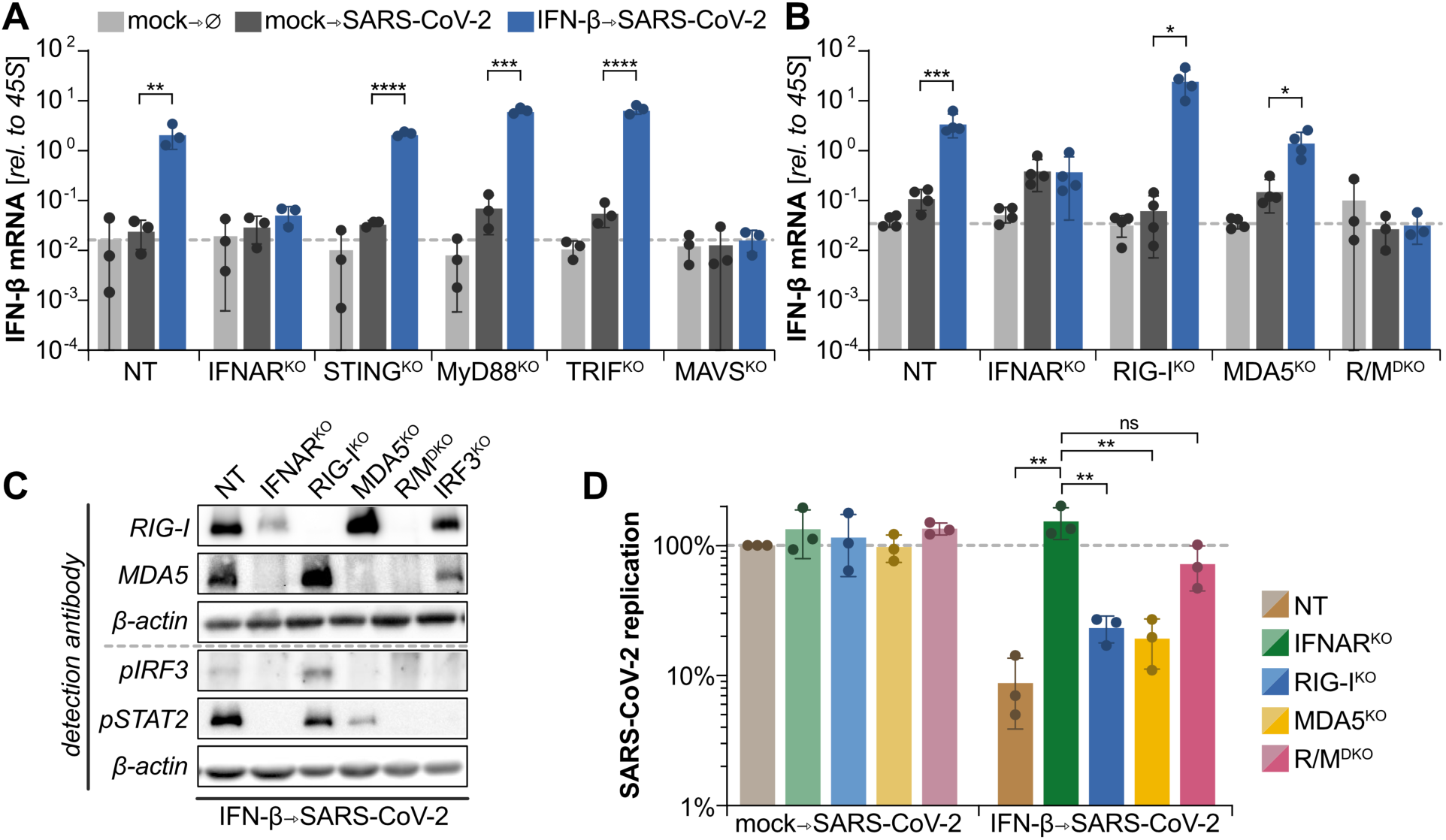
PRR-dependence of the antiviral response to SARS-CoV-2 in primed cells. A549^ACE2/TMPRSS2^ cells harboring the indicated CRISPR/Cas9-mediated functional KOs of **(A)** the major PRR adaptors or of **(B)** the RLRs (R/M^DKO^: RIG-I/MDA5 double-KO) were either mock-treated or primed with IFN-β (see Fig. 1) followed by infection with SARS-CoV-2 (MOI of 0.1). At 24 hours post infection, IFN-β (*IFNB1*) gene expression was determined by qRT-PCR. Values were normalized to 45S rRNA levels. Bar represent mean ±SD of three independent experiments (individual experiments shown as dots). NT: non-targeting CRISPR gRNA used as wildtype control; IFNAR^KO^ cells cannot be primed by IFN-β and were used as controls. **(C)** Primed and infected A549^ACE2/TMPRSS2^ cells (as in A, B above) with the indicated KOs were lysed 24 hours post infection and analyzed by immunoblotting using the indicated antibodies (“*p*” denotes phosphorylation specificity). Blot representative of two biologically independent experiments. **(D)** Mock-treated or primed A549^ACE2/TMPRSS2^ cells harboring the indicated functional KOs (see panel B) were infected with SARS-CoV-2 (MOI 0.1). At 24 hours post-infection, intracellular viral genome levels were determined by qRT-PCR using primers specific for ORF1. Values were normalized to 45S rRNA levels and expressed relative to non-primed wildtype (NT) cells. Bars represent mean ±SD of three independent experiments (individual experiments shown as dots). Statistical significance was tested by an unpaired two-tailed (**A, B**) and one-tailed (**D**) t-test; * p<0.05, ** p<0.01, *** p<0.001, and **** p<0.0001.

While our data clearly showed the restoration of an RLR-mediated IFN-response towards SARS-CoV-2 infection in IFN-β-primed cells, it remained unclear whether this response was effectively antiviral. Therefore, we next assessed viral replication upon infection of IFN-β-primed cells. We again used a low dose (1x IC50) of IFN-β for pre-treatment of A549^ACE2/TMPRSS2^ cells and then infected them with SARS-CoV-2. As expected, viral replication at 24 hours post infection was significantly (∼10-fold) reduced in IFN-pre-treated cells as compared to mock-treated cells (Figure 2D, brown bars), while no effect was observed in IFNAR^KO^ control cells lacking the type I IFN receptor (Fig. 2D, green bars). Strikingly, IFN-β-pre-treatment tended to be less effective in suppressing viral replication in cells lacking functional MDA5 or RIG-I, and almost completely ineffective (no statistically significant effect) in RIG-I/MDA5^DKO^ cells lacking both receptors (Fig. 2D, pink bars). This was surprising as generally the direct induction of ISGs downstream of IFN receptor signaling was held accountable for the antiviral effects of IFN-treatment. Our results now underscore the critical importance of RLR-mediated positive feedback for the effects of low-dose IFN. Indeed, at higher doses of IFN-β (10x IC50) the effects of pre-treatment were significantly stronger and virtually independent of the presence of RLR feedback (Suppl. Fig. 2C). These observations warrant the notion that low local concentrations of IFNs prime tissue cells for more sensitive detection of potential viral infection, rather than directly eliciting an antiviral state.

### Elevated levels of MDA5 or RIG-I suffice to induce IFN production upon SARS-CoV-2 infection

The above experiments suggested IFN-priming of cells increases the sensitivity of the antiviral system largely by inducing expression of the viral sensors. Nonetheless, IFN-treatment upregulates a plethora of genes, possibly also factors downstream of the RLRs. In order to assess if the enhanced expression of only RIG-I or MDA5 suffices to reconstitute productive viral sensing, we generated stably overexpressing cells. We reconstituted RIG-I or MDA5 expression in double KO cells (A549 RIG-I/MDA5^DKO^) by lentiviral transduction under the control of the weak ROSA26 promoter (Fig. 3A, bottom). This led to the expression of moderate amounts of either RIG-I or MDA5, without triggering signaling and IFN induction on its own (Fig. 3A, light bars). Upon infection with SARS-CoV-2, however, IFNB1 transcription was significantly induced both in RIG-I and MDA5 overexpressing cells (Fig. 3A, dark bars). These experiments demonstrate that, indeed, the enhanced expression of RLRs suffices to enable cells to mount an IFN response towards SARS-CoV-2 despite fast and strong viral antagonism. This corroborates our previous observation that higher expression of viral sensors in the nasal airways of children prior to infection associates with significantly stronger IFN responses upon SARS-CoV-2 infection (20). Briefly, in that earlier study we analyzed nasal airway swabs of healthy and of SARS-SoV-2-infected children and adults by scRNA-Seq. Now focusing exclusively on the samples of healthy donors, comprising 18 children (4-16 years, median age 9.0) and 23 adults (24-77 years, median 46.0), finer analysis of the data revealed a gradual decrease of MDA5 expression levels in nasal epithelial cells with increasing age of the individuals (Fig. 3B). RIG-I was also confirmed to be significantly higher expressed in the epithelium of children and adolescents, however, the gradual decrease between age groups was less clear (Fig. 3B).

**Figure 3.**
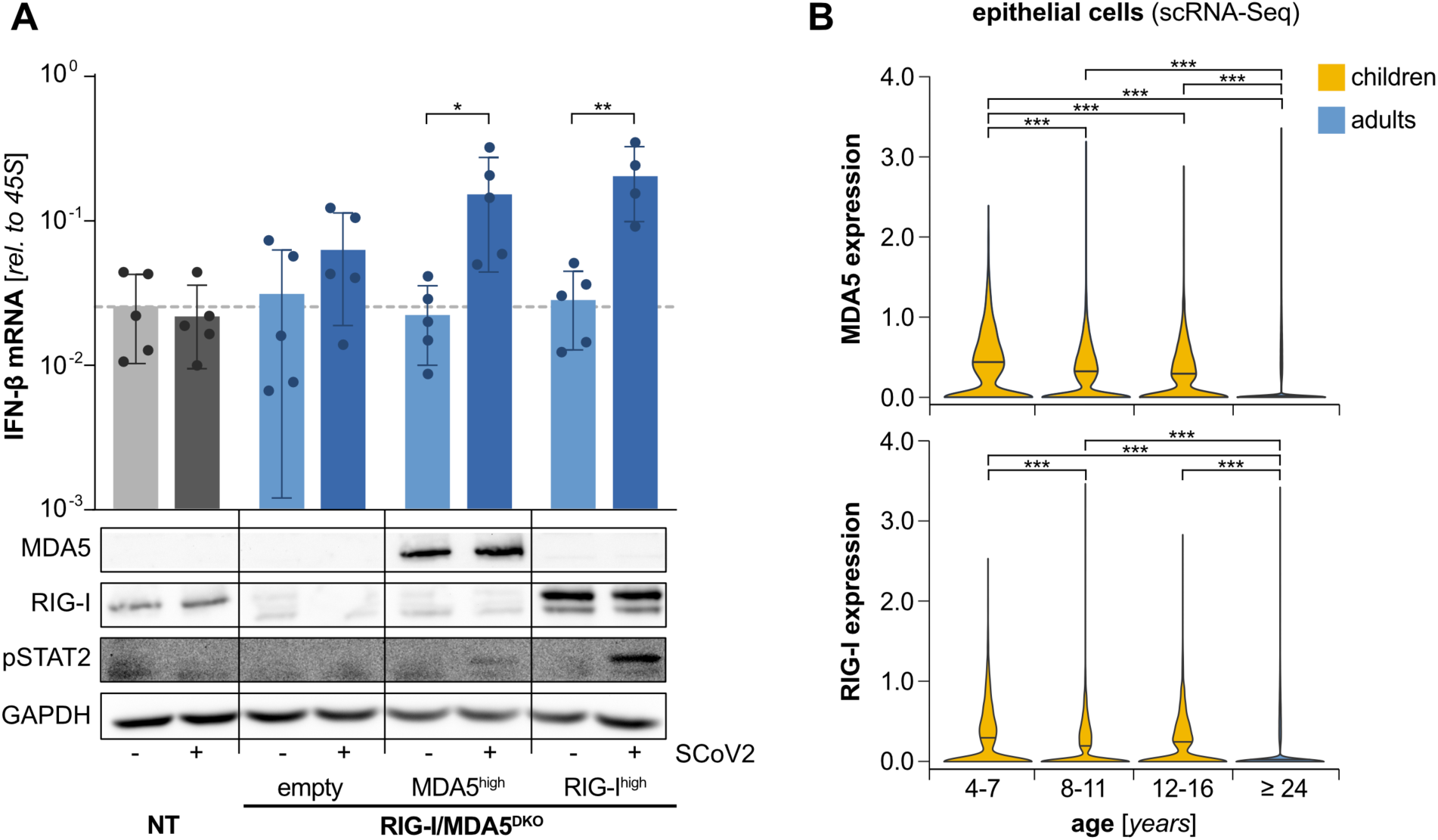
Impact of increased RLR expression on antiviral response to SARS-CoV-2 in non-primed cells. **(A)** A549^ACE2/TMPRSS2^ cells deficient for RIG-I and MDA5 (DKO: double knockout) were reconstituted to express only MDA5 or RIG-I by lentiviral transduction (under control of the weak ROSA26 promoter). Cells were mock-infected or infected with SARS-CoV-2 (upper panel: MOI 0.01, lower panel: MOI 1) for 24 h. IFN-β (*IFNB1*) expression was determined by qRT-PCR (bar graph), and MDA5 and RIG-I protein levels as well as STAT2 phosphorylation (pSTAT2) were analyzed by immunoblotting (lower panel). qPCR values were normalized to 45S rRNA and expressed as mean ±SD of at least four independent biological experiments (individual experiments shown as dots). Statistical significance was tested by a paired two-tailed t-test. * p<0.05 and ** p<0.01. Immunoblot is representative of two independent experiments. **(B)** Nasal swabs were taken from heathy individuals of the indicated age groups and analyzed by scRNA-Seq (20). MDA5 (*IFIH1*) and RIG-I (*RIGI*) gene expression were quantified across all epithelial cell types and displayed as violin plots (median indicated). Statistical significance was tested using two-tailed Wilcoxon comparison adjusted with Bonferroni correction. *** p < 0.001.

A growing body of evidence established a rapid and strong IFN response locally in the infected tissue as one of the most important characteristics of a successful immune clearance of SARS-CoV-2 and, hence, the prevention of severe manifestations of COVID-19 (9, 10, 19, 51). We have previously described the importance of a primed state of epithelial cells for the mounting of a swift and robust IFN response upon SARS-CoV-2 infection in children (20). We now showed the critical role of *a priori* expression levels of the viral sensors RIG-I and MDA5 in this epithelial cell priming and found RLRs to be particularly highly expressed in children and teenagers.

### Immune cells stimulate RLR expression in epithelial cells in an IFN-dependent and-independent manner

We now aimed to better understand the background of the increased RLR expression in children. The homeostatic set-point of cell-intrinsic immunity of epithelial tissue is determined by the interplay of epithelial cells and tissue resident and other local immune cells (52, 53). Therefore, we again turned to the scRNA-Seq data of nasal swabs of healthy children and adults. We found vastly higher proportions of immune cells in the nasal mucosa of children (>50%) than that of adults (<10%), in particular neutrophils, T-cells, macrophages and dendritic cells (Figure 4A, finer subtyping of cells in Suppl. Fig. 3A) (20). We then analyzed immune–epithelial cell interactions by mapping of ligand-receptor-pair expression at the single cell level onto the CellChat database (41), augmented with a recently published very comprehensive interaction map of the human immune system (42). Strikingly, we observed substantially more but also stronger inferred immune– epithelial cell interactions in children than in adults (Figure 4B, see also Suppl. Fig. 3B-D). We speculate that the increased presence of immune cells in the non-sterile environment of the nasal mucosa may shape the local cytokine milieu and is decisive for the heightened expression of RIG-I and MDA5. In order to approach this in a simplified *in vitro* experimental model, we used isolated human immune cells (peripheral blood mononuclear cells, PBMC) and mimicked microbial encounter by exposing them to a Gram-negative model bacterium (*Yersinia enterocolitica, Ye*) (Fig. 4C). To prevent bacterial overgrowth, bacteria were killed after one hour by gentamicin. In order to functionally assess the ensuing cytokine response, we transferred the sterile-filtered PBMC supernatant to A549 cells 24 hours later. We observed a substantial increase in expression of both RLRs in A549 cells exposed to the supernatants of stimulated PBMC (Fig. 4D). As RIG-I and MDA5 are ISGs, this induction was likely to be mediated by IFNs produced by innate immune cells. Indeed, KO of the type I IFN receptor (IFNAR^KO^) in A549 cells dampened RLR induction significantly, and the additional KO of type II and III IFN receptors (IFNR^TKO^) further reduced it (Fig. 4D). Unexpectedly, however, even in the absence of all three classes of IFN receptors, both RIG-I and MDA5 levels were still significantly induced. We, therefore, analyzed the PBMC supernatants for their cytokine content. As expected, *Ye*-exposed PBMC secreted substantial amounts of type I (-α2, -β) and II (-γ) IFNs (Fig. 4E), coherent with the reduced RLR induction in IFN receptor KO cells. We further detected secretion of several pro-and anti-inflammatory cytokines, in particular IL-1β, IL-6, IL-10 and TNF, some or all of which may contribute to the observed RLR induction (Fig. 4E). In order to compare these *in vitro* findings to the physiological situation, we specifically analyzed the immune cell population from the nasal mucosa of healthy children and adults for their expression of these cytokines, incl. IFN-γ as a common inflammatory mediator. Strikingly, we found significantly higher expression for IFN-γ, IL-6, IL-10 and TNF (but not IL-1β) in healthy children as compared to adults (Fig. 4F), resembling and confirming our *in vitro* observations. To further understand the source of these cytokines and their contribution to immune–epithelial communication *in vivo*, we analyzed ligand–receptor interactions of the single cell data specifically for IFN-γ and TNF (Fig. 4G) as well as IL-1 and IL-10 (Suppl. Fig 3 E-G); there was no significant ligand–receptor interaction of IL-6. Indeed, all four analyzed cytokines exhibited notably higher connectivity to the epithelial cell compartment, both in numbers of interactions as well as the respective information flow (i.e. interaction strength). Interestingly, IL17A^+^ T-cells appeared to be major producers of IFN-γ, TNF and IL-10 in children, as well as proliferating NK cells for IFN-γ, and macrophages for IL-1 (Fig. 4G and Suppl. Fig. 3E).

**Figure 4.**
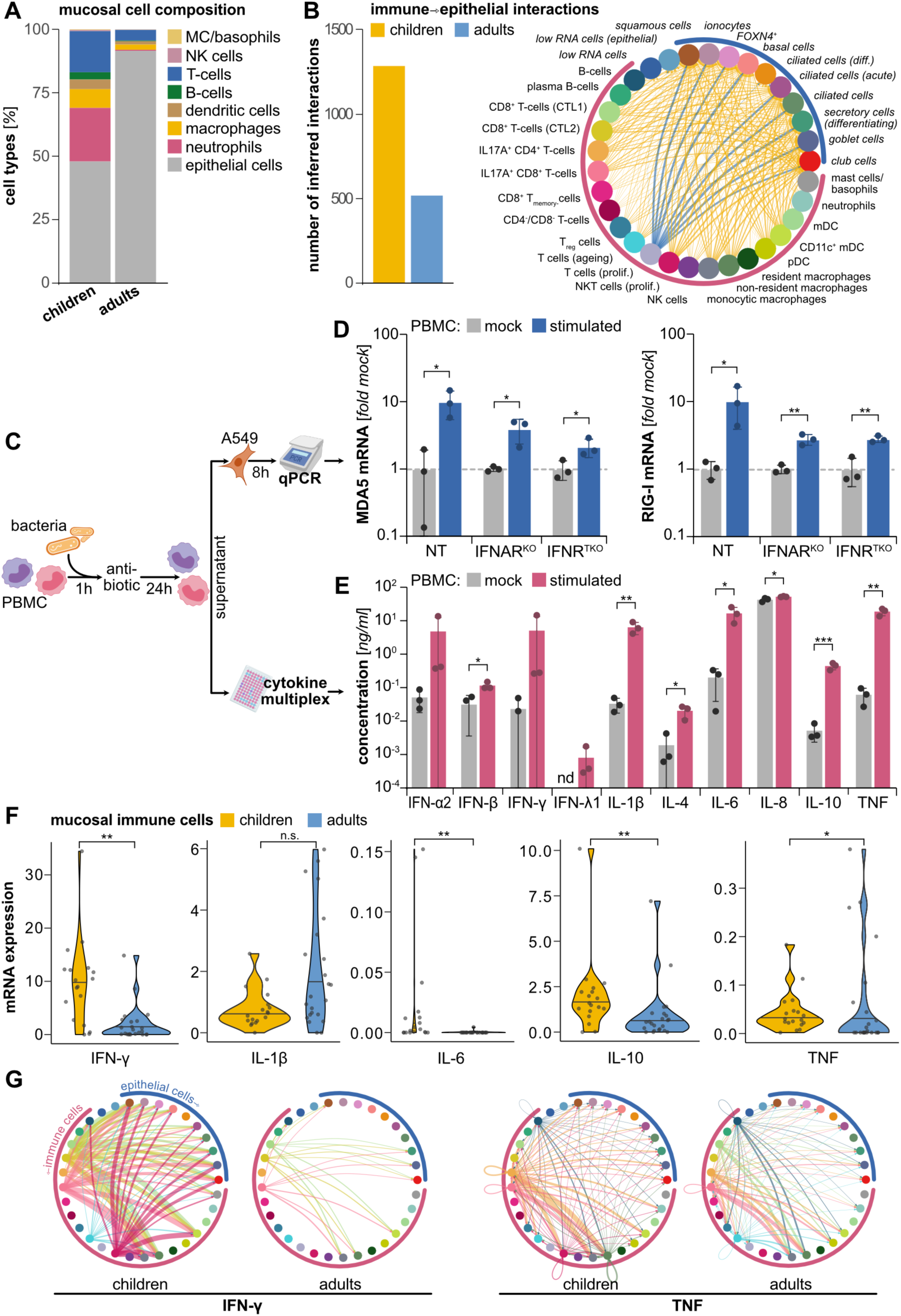
Antiviral priming of epithelial cells by immune – epithelial cell crosstalk through cytokines. Nasal swabs were taken from heathy children (4 – 16 years, n=18) and adults (≥24 years, n=23) and analyzed by scRNA-Seq (previously published in (20)). **(A)** Cell types and states were assigned to single cells based on their transcriptional profile (for details see (20)), and similar types were subsumed under more general classes (e.g. “epithelial cells”, “T-cells”). The relative abundance of immune cell types were determined and compared between children and adults. Detailed immune cell composition with individual cell types can be found in Suppl. Fig. 3. **(B)** Based on expression of corresponding pairs of signaling ligands (on immune cells) and receptors (on epithelial cells), the interactivity of mucosal immune and epithelial cells was determined in children and adults. Total numbers of inferred interactions are shown in the bar graph (left). Interactions of specific immune cell types with epithelial cell types are shown in the interaction plot (right). The number of underlying ligand-receptor-interactions is coded by line thickness, with higher numbers in children shown in yellow and those higher in adults shown in blue. The circle segments around the plot signify cell class (blue: epithelial, pink: immune cells). See also Suppl. Fig. 3. **(C)** Isolated human peripheral blood mononuclear cells (PBMC) were exposed to live Gram-negative bacteria (*Yersinia enterocolitica, Ye*) for 1 hour, after which bacteria were killed by antibiotic (gentamicin) treatment. Upon further incubation for 24 hours, culture supernatants were harvested and analyzed. **(D)** A549 NT (wildtype control), IFNAR1^KO^ and type I, II, III IFN receptor triple KO (IFNR^TKO^) were incubated with collected PBMC supernatants for 8 hours. Expression levels of MDA5 (*IFIH1*) and RIG-I (*RIGI*) were determined by qRT-PCR. Values were normalized to GAPDH levels and expressed relative to wildtype A549 incubated with supernatants from mock-stimulated PBMC. Bars represent means ± SD of three biologically independent experiments (individual experiments shown as dots). **(E)** Supernatants of mock-treated PBMC or of PBMC exposed to *Ye* were analyzed for their cytokine content by electroluminescent multiplex assays (MSD platform). Data is shown as means ± SD from three independent experiments (individual experiments shown as dots). **(D, E)** Statistical significance was tested by an unpaired one-tailed t-test. * p < 0.05, * p < 0.01, ** p < 0.01, and *** p <0.001. **(F)** ScRNA-Seq data from nasal swab material of healthy children and adults (as in A) were analyzed for the expression of cytokines (see E) across all immune cell types and states. Values displayed as violin plots with average per cell expression level of each donor shown as dots. Statistical significance was tested using Kolmogorov–Smirnov test with Benjamini-Hochberg correction. **p<0.01, *p<0.05. **(G)** ScRNA-Seq data were analyzed for ligand – receptor interactions between cell types (cell type identified by color, see panel B) specifically for IFN-γ (left) and TNF (right) signaling in children and adult samples. Interaction strength is coded by line thickness, direction (ligand⇾receptor) indicated by arrowheads, arrows are color-coded by cell type of origin. See also Suppl. Fig. 3.

In summary, we found that expression of RIG-I and MDA5 in epithelial cells can be stimulated by inflammatory mediators released by activated immune cells. The nasal epithelium of children contains a substantially higher number of such activated immune cells that show enhanced cytokine-based interaction with the epithelial cell compartment. In particular IFN-γ, TNF, IL-1β, IL-6 and IL-10 were identified upon PBMC stimulation *in vitro* and were further found to be elevated (at the mRNA level) in mucosal immune cells of children and/or to mediate a higher degree of interaction between immune and epithelial cells.

### IFN-independent priming induces MDA5 and RIG-I expression sufficiently to respond to SARS-CoV-2

This finding of enhanced immune–epithelial cross-talk in the mucosa of children was in line with our notion that constitutive presence of higher levels of certain cytokines would be responsible for the elevated basal expression of MDA5 and RIG-I in children. To further test which (if any) of the above-identified cytokines have the potential to induce RLR expression, we treated A549 lung epithelial cells with recombinant IFN-γ, IL-1β, IL-6, IL-10, or TNF at a concentration of 10 ng/ml, comparable to the amount measured in the supernatants of activated PBMC (compare Fig. 4E). Eight hours after stimulation, we measured MDA5 and RIG-I levels by qRT-PCR and found particularly IFN-γ and TNF, and to a lesser extent IL-1β, to induce MDA5 and RIG-I transcription (Fig. 5A). As to be expected, IFN-γ required presence of the IFN-γ-receptor (IFNGR) and RLR induction was abolished in IFNR^TKO^ (but not IFNAR^KO^) cells. In contrast, the effects of TNF and IL-1β were fully independent of any IFN-signaling (Fig. 5A). IL-6 and IL-10 did not affect RLR expression in A549 cells at all.

**Figure 5.**
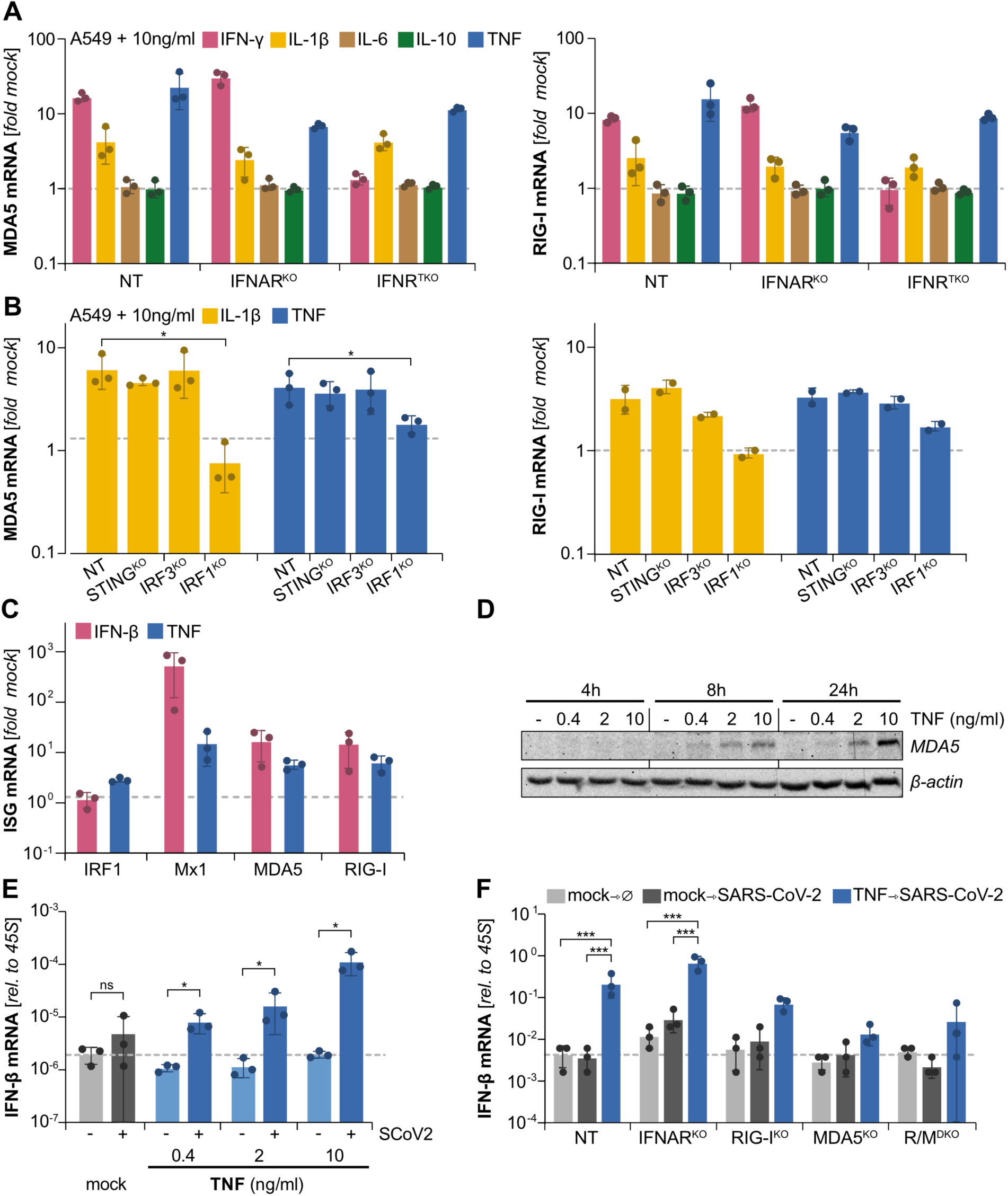
Epithelial cell priming by the proinflammatory cytokine TNF. **(A)** A549 cells, wildtype (NT, non-targeting control) or with the indicated CRISPR/Cas9-mediated functional KO, were treated with 10 ng/ml of individual recombinant human cytokines as indicated. After 8 hours, cells were lysed and expression of MDA5 (*IFIH1*) and RIG-I (*RIGI*) was determined by qRT-PCR. Values were normalized to GAPDH and expressed relative to mock-treated cells. **(B)** A549 cells, wildtype (NT) or with the indicated KO, were treated with 10 ng/ml IL-1β or TNF, and MDA5 (*IFIH1*) and RIG-I (*RIGI*) expression was determined as in (A). **(C)** A549 cells were treated with TNF (10 ng/ml) or with IFN-β (1x IC50, 170 IU/ml) for 16 hours. Expression of the indicated IFN stimulated genes (ISG) was determined by qRT-PCR. Values were normalized to GAPDH and expressed relative to mock-treated cells. **(D)** A549 cells were treated with recombinant human TNF at the indicated concentrations (or mock-treated) for 4, 8, or 24 hours. Protein expression of MDA5 was determined by immunoblotting (β-actin shown as loading control). Blot is representative of two independent experiments. **(E)** A549^ACE2/TMPRSS2^ cells were mock-treated or primed with TNF at the indicated concentrations for 8 hours followed by infection with SARS-CoV-2 (MOI 1). At 24 hours post-infection, cells were lysed and induction of IFN-β (*IFNB1*) gene expression was determined by qRT-PCR. Values were normalized to 45S rRNA. **(F)** A549^ACE2/TMPRSS2^ cells with no (NT) or the indicated KO (R/M^DKO^: RIG-I and MDA5 double KO) were mock-treated or primed with TNF (10 ng/ml) for 8 hours followed by SARS-CoV-2 infection (MOI 1). At 24 hours post infection, induction of IFN-β (*IFNB1*) expression was determined by qRT-PCR. Values were normalized to 45S rRNA. **(A, B, C, E, F)** Bars represent the mean ±SD of three independent experiments (individual experiments shown as dots). **(B, E, F)** Statistical significance was tested by an unpaired one-tailed t-test. * p < 0.05, *** p <0.001 and **** p < 0.0001.

We next investigated how these cytokines might upregulate RLRs, which clearly are not amongst their canonical target genes. TNF and IL-1β are hallmark pro-inflammatory cytokines. Recently, they were described to be capable of indirectly inducing ISG expression via the release of mitochondrial DNA, which then activates the cytosolic cGAS-STING-IRF3 pathway leading to the production of type I IFN (54, 55). While secreted IFN was dispensable for RLR induction by TNF and IL1-β in our experiments (Fig. 4G), direct induction downstream of cGAS and STING was still plausible. Therefore, we checked MDA5 and RIG-I expression in TNF-or IL-1β-stimulated STING^KO^ and IRF3^KO^ cells. Surprisingly, at 8 hours post-stimulation, both STING and IRF3 deficient cells induced RLR transcription to comparable levels as WT cells, suggesting that early RLR induction occurred independently of the cGAS-STING axis (Fig. 5B). Another important transcription factor of the IRF family is IRF1. Recently, IRF1 was described to be involved in constitutive expression of antiviral genes, such as TLR2, TLR3, and OSA2 (56), and to contribute to TNF-and IL-1β-mediated ISG expression in monocytes and epithelial cells (55, 57). In fact, in IRF1^KO^ cells we observed virtually no induction of RIG-I and MDA5 transcripts upon IL-1β treatment, and also TNF-induced expression was clearly reduced (Fig. 5B). Notably, IRF1 is also an important and well-established transcription factor in the IFN-γ response (58–60). We furthermore found elevated IRF1 expression in almost all epithelial and many immune cell types in the nasal mucosa of children as compared to adults (Suppl. Fig 4A). Taken together, from the cytokines identified *in vitro* and *in vivo* above, we found TNF, IFN-γ and, to a lesser extent, IL-1β to be capable of stimulating the expression of RIG-I and MDA5 in A549 epithelial cells. This suggests those cytokines critically determine homeostatic expression of the antiviral system in the epithelium, particularly of children, with IRF1 being the central transcriptional regulator.

Both, *in vitro* and *in vivo* IL-1β appeared to play a minor role. IFN-γ is antiviral and its signaling and transcriptional response has significant overlap with type I/III IFN, hence, its function in priming of cells likely resembles IFN-β (Fig. 1, 2). TNF, on the other hand, is a highly pleiotropic cytokine involved in organogenesis, tissue homeostasis and, importantly, it is a critical mediator of inflammatory responses. Its biological activity is highly context– and likely concentration– dependent. We therefore further investigated the capacity of IFN-independent priming of epithelial cells by TNF. Confirming our above finding of IRF1-dependence, TNF-treatment induced transcription of IRF1 to higher levels than IFN-β, whereas the classical type I ISG Mx1 was substantially higher in IFN-treatment (Fig. 5C). The RLRs MDA5 and RIG-I, however, were induced similarly by both cytokines. We next assessed MDA5 expression at the protein level upon different TNF doses and treatment times. Indeed, protein levels increased in a dose-and time-dependent manner (Figure 5D). We then pre-treated A549^ACE2/TMPRSS2^ cells with increasing TNF doses for 8 hours, before infecting them with SARS-CoV-2. Twenty-four hours post infection we then analyzed the ensuing antiviral response by measuring IFN-β mRNA levels. While TNF pre-treatment alone did not induce IFN-β, SARS-CoV-2 infection of TNF-primed cells in fact induced significant levels of IFN-β mRNA in a dose-dependent manner (Figure 5E). In order to corroborate the central role of viral sensing through the RLRs for IFN-induction also in TNF-primed cells, we stimulated RLR deficient cells (RIG-I or MDA5 individual KO, or double KO) and checked for IFN-β production upon infection. Depletion of RIG-I slightly reduced IFN-β production at the mRNA (Fig. 5F) and protein level (Suppl. Fig. 4B), whereas IFN-β was almost completely diminished in MDA5^KO^ and RIG-I/MDA5^DKO^ cells (Fig. 5F, Suppl. Fig. 4B). This effect was confirmed for the induction of the ISG RSAD2 (Suppl. Fig. 4C). As expected, in contrast to IFN, TNF-pre-treatment was not impacted by IFNAR deficiency, corroborating TNF-priming is independent of type I IFN signaling (compare Fig. 4). It further demonstrates that IFN-β induction in TNF-primed cells does not require positive feedback through IFN signaling. Interestingly, IFNAR^KO^ cells even tended to exhibit a slight increase in sensitivity, but below statistical significance (Fig. 5F, Suppl. Fig. 4).

Overall, these data suggest that similar to type I IFN, TNF can prime the antiviral sensing machinery of epithelial cells sufficiently to restore IFN-induction upon SARS-CoV-2 infection. This sensitization is largely mediated by MDA5, likely through its strong induction by TNF signaling.

### Immune cells from children are more effective in inducing MDA5 and mounting an antiviral defense against SARS-CoV-2

So far, our experiments showed that immune cells via cytokine release can tune the antiviral responsiveness of epithelial cells, largely through regulating the expression levels of viral sensors such as RIG-I and MDA5. We observed this to be particularly prominent in the airway epithelium of children, correlating with their significantly stronger induction of antiviral responses upon SARS-CoV-2 infection as compared to adults (20). We now wanted to understand whether the increased number of immune cells in the mucosal tissue of children (Fig. 4A) alone explains the stronger priming of epithelial cells, or whether immune cells of younger individuals additionally have a higher intrinsic potential to produce cytokines and stimulate RLR expression. We obtained PBMC from five healthy adolescents (ages 14-17, median: 15.5 years) and six adults (ages 35-66, median: 46,25 years); note these individuals were not participants in our previous scRNA-Seq study (20). We again stimulated the PBMC by one hour exposure to live *Ye*, after which we killed the bacteria by gentamicin (Fig. 6A). Twenty-four hours later, supernatants of PBMC were transferred to A549 cells and MDA5 and RIG-I mRNA levels were assessed after 8 hours of incubation. While the supernatants of non-stimulated PBMC had no impact on RLR expression, supernatants of bacteria exposed immune cells strongly induced both MDA5 and RIG-I mRNA (Fig. 6B). Interestingly, PBMC supernatants of the younger individuals elicited a significantly higher MDA5 expression in A549 cells than those of adults. RIG-I expression showed a similar but non-significant trend. When we assessed the IFN content of PBMC supernatants, we observed slightly increased protein levels of all measured IFNs (IFN-α2, -β, -γ, and -λ1) in the PBMC from adolescents as compared to those of adults, albeit none reached statistical significance (Figure 6C, further cytokines see Suppl. Fig. 5). These findings indeed argue in favor of an intrinsically more vigorous response of immune cells of younger individuals than cells of adults towards an identical stimulus.

**Figure 6.**
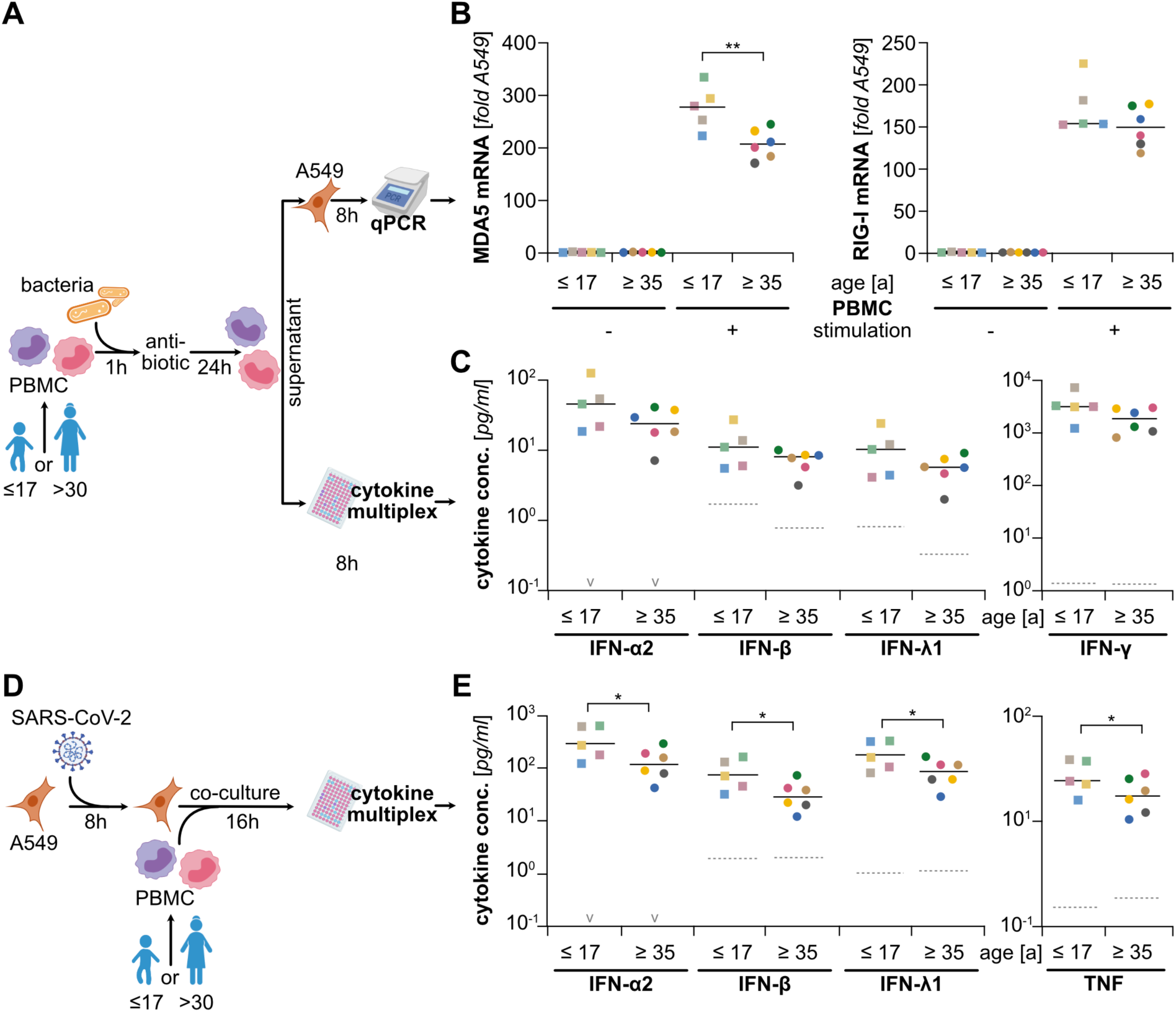
Antiviral priming of epithelial cells by immune cells of juvenile *versus* adult donors. **(A)** Peripheral blood mononuclear cells (PBMC) isolated from healthy juvenile (14 – 17 years) and adult (35 – 60 years) donors were mock treated or exposed to live Gram-negative bacteria (*Yersinia enterocolitica, Ye*) for 1 hour, after which bacteria were killed by antibiotic (gentamicin) treatment. Upon further incubation for 24 hours, culture supernatants were harvested and analyzed. **(B)** A549 cells were incubated with collected supernatants from mock treated or bacteria-stimulated PBMC. After 8 hours, expression of MDA5 (*IFIH1*) and RIG-I (*RIGI*) were determined by qRT-PCR. Values were normalized to GAPDH levels and expressed relative to A549 not treated with PBMC supernatant. **(C)** Supernatants of mock-treated PBMC or of PBMC exposed to *Ye* were analyzed for their IFN content by electroluminescent multiplex assays (MSD platform). **(D)** A549 cells were infected with SARS-CoV-2 (MOI 0.1) for 8 hours before adding PBMC to the culture. After 16 hours of co-culturing, supernatants were harvested and subjected to cytokine multiplex measurement. **(E)** Levels of secreted cytokines were determined (see Suppl. Fig. 6B for further cytokines). **(B, C, E)** Values for each donor are shown individually (5 juvenile, 6 adult donors), color identifies donor across all measurements. Median shown as solid black line; median of (B, C) mock stimulated samples or (E) co-culture of uninfected A549 with PBMC shown as dashed gray line (individual data points omitted), “v” signifies median below 10^-1^ pg/ml. Statistical significance was tested by an unpaired one-tailed t-test. *p < 0.05, ** p < 0.01.

Lastly, having observed this stronger response of juveniles’ immune cells, we were curious whether this propensity would also lead to more vigorous responses to SARS-CoV-2 infection. It has previously been reported that co-cultivation of infected epithelial cells with plasmacytoid dendritic cells (pDC) allows sensing of infection and production of type I IFN by pDC (61). We aimed at reproducing this by co-culturing SARS-CoV-2 infected A549^ACE2/TMPRSS2^ cells with PBMC from the same adolescent and adult donors as above. We added PBMC to the epithelial cells 8 hours after SARS-CoV-2 infection. After 16 hours of co-cultivation, we then determined the secreted cytokine profile (Fig. 6D). While in the absence of PBMC, A549 cells once more failed to secrete appreciable amounts of IFNs (Supp. Fig. 6A), we measured substantial amounts of type I, II and III IFNs as well as TNF in the co-culture setting (Fig. 6E, Supp. Fig. 6A). Also in this set-up, we found a statistically significant tendency of higher IFN and TNF levels in the co-cultures with PBMC of adolescents as compared to adults (Fig. 6E). Interestingly, this cytokine production by immune cells likely in turn sensitized virus sensing and IFN-β production by the A549 cells, as co-culture of PBMC with infected RLR-deficient (RIG-I/MDA5^DKO^) A549 cells yielded significantly lower IFN-β and IFN-λ1 levels (Supp. Fig. 7).

Overall, our data indicate that in children and adolescents epithelial cells of the nasal mucosa are potently primed due to at least two distinct factors: 1) a higher density of immune cells in the mucosal tissue, and 2) a more vigorous cytokine production by those immune cells. The latter most likely also contributes to a faster and stronger mounting of an antiviral response to SARS-CoV-2 infection. In effect, this likely forms the basis for the more efficient immunological control of the infection in children and the substantially lower risk of pediatric patients developing severe courses of COVID-19.

## Discussion

A strong and well-coordinated immune response is key for the successful clearance of any viral infection. A central role in the coordination of the many effector branches of the immune system is assumed by early and little-specific innate immune responses (62). Given the extreme replication kinetics not uncommon with viruses, a swift and potent induction of type I and III IFNs– the major antiviral mediators– by the infected cells themselves as well as, secondarily, local tissue-patrolling innate immune cells is key in proper and timely triggering of a full-fledged and efficacious defense program. The swiftness of virus recognition and the onset of IFN production is even more important given many viruses quickly express potent antagonists blunting specifically these immediate-early host defense pathways (63). This race-like situation is infamously exemplified by the recently emerged SARS-CoV-2 combining very fast replication kinetics (17) with a versatile array of immune antagonists that effectively interfere with many steps of the cell-intrinsic antiviral system (reviewed in (25)). As a consequence, SARS-CoV-2 infection fails to induce significant amounts of type I and III IFNs in immune competent A549 lung epithelial cells (18, 23) as we also confirm in the present study, as well as in further cell culture models. Nonetheless, some reports describe notable production of IFNs in certain cells, including lung and intestinal epithelial cells (64–66). We argue that this apparent contradiction is a direct consequence of the race-like virus-host interplay early in infection, which is exceptionally dichotomized by SARS-CoV-2: either the cell senses and responds to the virus fast enough, or the virus shuts off the respective pathways virtually completely. Indeed, we here demonstrate that SARS-CoV-2 non-responsive A549 cells can be rendered fully responsive by pre-treating them with low-moderate doses of IFN or TNF, underscoring that subtle differences in the state of the host cell can be the basis of an all-or-nothing outcome in terms of IFN production.

The efficient suppression of IFN induction by SARS-CoV-2 appears to be a major determinant for viral replication and spread and, hence, for disease development. Even though viral antagonists also target IFN downstream signaling (25), overall the virus remains very sensitive to IFN (67–69). In fact, clinical genetics studies clearly established a malfunctioning type I IFN response as a critical determinant predisposing individuals for severe and life-threatening COVID-19 (9, 10), however with incomplete penetrance particularly in adolescents and young adults (70). While high IFN levels in the lung and serum at later stages of the disease associate with severity (2, 12, 13), early and local IFN production right at the primary infection site in the upper respiratory tract has been appreciated as a major determinant for successful control of infection and mild courses of disease (11, 14, 15, 17, 51). By single cell analyses of cells isolated from nasal swabs we and others could show that the high resilience of children towards severe COVID-19 is associated with an increased expression of virus sensors, in particular MDA5, in airway epithelial cells already prior to infection and, consequently, with a significantly faster and stronger IFN response upon SARS-CoV-2 infection as compared to adult patients (19, 20).

Here we investigated the molecular basis of efficient detection of SARS-CoV-2 infection and the mounting of a proper IFN response. For this purpose, we have set up a simplified *in vitro* model based on the lung epithelial cell line A549, and validated our findings in scRNA-Seq data from nasal swab samples. A549 cells are well-known to have a functional and potent antiviral system (34, 39), but are little responsive to SARS-CoV-2 infection, similar to what we have previously observed for airway epithelial cells of adult donors (20). In that study we found the primed state of epithelial cells in children to be somewhat reminiscent of an IFN signature that included the increased expression of the RLRs. We now mimicked this state *in vitro* by pre-treatment of A549 cells with a rather low dose of IFN-β. This led to the increased expression of a variety of pattern recognition receptors, allowing us to study their respective impact on functional SARS-CoV-2 recognition. It has previously been shown that MDA5 is the major receptor sensing SARS-CoV-2 in epithelial cells (26–28, 48). Employing an array of CRISPR/Cas9-based functional KOs, we confirmed MDA5 as the single most important sensor in this system. Notably, there was no impact of TLR/TRIF/MyD88 or cGAS/STING pathways whatsoever in our system. However, we identified a significant contribution by RIG-I: only simultaneous depletion of both receptors, MDA5 *and* RIG-I, completely abrogated IFN-β production upon SARS-CoV-2 infection. Correspondingly, moderately elevated expression of RIG-I only– in the complete absence of MDA5– sufficed to sense infection and trigger the IFN-β response. The role of RIG-I in sensing of the virus has not been widely appreciated before, but was shown in one previous study (47). Moreover, RIG-I has been implicated with IFN-and signaling-independent inhibition of SARS-CoV-2 replication by direct binding to the 3’-UTR of the viral genome (71). In fact, in RIG-I^KO^ cells, we even observed a slight increase in IRF3 phosphorylation and ensuing IFN-β, which might be related to somewhat stronger viral replication (not confirmed in qRT-PCR) or increased accessibility to MDA5 in the absence of RIG-I binding.

The observed IFN-like signature in the epithelium of children as well as our IFN-β priming approach both are associated with the induction of countless genes. However, our KO studies in A549 cells confirmed the crucial role of the RLRs for the detection of SARS-CoV-2. Furthermore, our complementary approach expressing only either MDA5 or RIG-I on a double-KO background to levels slightly above the endogenous situation was sufficient to render cells responsive to SARS-CoV-2 infection similar to IFN pre-treatment. This supports the notion that elevated expression specifically of RLRs would be causal for a stronger IFN production in the upper airways of children as compared to adults. It is further supported by the fact that particularly MDA5 expression is highest in infants and gradually decrease over childhood and adolescence, mirroring the increasing risk for severe disease with age (6). So far, it is unclear whether a further decrease in RLR expression may be responsible also for the exponentially increasing risk for severe COVID-19 in the elderly. A previous study showing reduced vigor of RIG-I signaling in monocytes of aged individuals as one aspect of immunosenescence may support this idea (72).

This age-dependence of the antiviral responsiveness of epithelial cells may be due to a developmental program. In fact, it has been shown that a subset of ISGs is highly expressed in the stem cell compartment and is lost during differentiation (73). However, this stem cell program appears to be tuned for direct antiviral effectors at the expense of the inducible IFN response, and indeed, neither RIG-I nor MDA5 are expressed to significant levels in stem cells (73). Alternatively, the elevated expression in airway epithelia of infants and school-age children may be mediated by immune–epithelial cell interactions (52, 53). Interestingly, we find substantially higher proportions of immune cells in the mucosa of healthy, i.e. non-infected, children as compared to adult donors (see also (20)) and we also find stronger immune–epithelial cell interactivity. We hypothesize immune cells, particularly upon frequent microbial encounter in a non-sterile environment such as the upper respiratory tract, give rise to a cytokine milieu capable of priming the antiviral sensing machinery of epithelial cells. This hypothesis was confirmed *in vitro* where we found bacteria-exposed immune cells to potently stimulate RLR expression in epithelial A549 cells through cytokines such as IFN-α, IFN-γ, TNF and to some lesser extent IL-1β. Indeed, also *in vivo* we found IFN-γ and TNF to be significantly higher expressed in mucosal immune cells of healthy children than in those of adults. Interestingly, we did not find significant expression of type I or III IFN in our scRNA-Seq data. This may be due to technical reasons, but in the organism those IFNs are also stringently restricted to the acute phase of an infection in order to prevent harmful side effects (74). Importantly, despite RIG-I and MDA5 being classical ISGs, type I IFNs were not decisive in epithelial RLR upregulation by immune cells, and even in fully IFN-blind (type I, II and III) A549 IFNR^TKO^ cells the RLRs were still significantly induced. This is in line with other studies in which MDA5 could be induced in an IFN-independent manner, e.g. by TNF (57, 75). Although recently both IL-1β and TNF were shown to lead to mitochondrial DNA release and signaling through the cGAS/STING/IRF3 pathway to induce IFNs and ISGs (55), in KO experiments we could show that in our system RLR upregulation was completely independent of STING and IRF3, but instead almost entirely dependent on IRF1. While this was surprising at first, it is in line with reports ascribing a central role to IRF1 in regulating constitutive expression of antiviral genes, such as TLR2, TLR3, and OAS2 (56), and contributing to TNF-and IL-1β-mediated ISG expression in monocytes and epithelial cells (55, 57). Interestingly, IRF1 is also a major transcription factor downstream of IFN-γ, making it a central regulator for the antiviral priming of airway epithelial cells. Taken together, this suggests immune cells in the respiratory mucosa tune the antiviral system of epithelial cells through creating a weak but constitutive inflammatory cytokine milieu. It remains to be elucidated whether an increased frequency of microbial encounter by children, e.g. during play or generally less hygienic behavior, contributes to immune cell activation. Importantly, though, even upon experimental stimulation with equal amounts of bacteria, we observed a stronger potential of PBMC of young donors to induce MDA5 and RIG-I expression in A549 cells as compared to adult PBMC, presumably due to a slight tendency for increased cytokine production. This once more highlights a higher reactivity of the (innate) immune system of younger individuals. The opposite effect has been described as immunosenescence, referring to less potent immune responses in the elderly (76), and might very well be a major contributor to the observed drastic increase in morbidity and mortality of COVID-19 in the aged population. Indeed, at the time of writing of this manuscript a very elegant study reported on the molecular underpinnings of exacerbated disease in old mice (77). The authors identified a lack of IFN-γ in aged mice to be the driving factor behind excessive viral replication and lack of immunological control of infection. Pre-treatment of mice with exogeneous IFN-γ almost completely reverted the aged phenotype. This appears to be a perfect mirror image of our above findings of IFN-γ (and TNF) being constitutively expressed at higher levels in the airways of young individuals, leading to antiviral priming of epithelial cells and a more potent overall immune response to SARS-CoV-2 infection.

Interestingly, while *in vitro* A549 cells did not produce notable amounts of IFNs and other cytokines in response to SARS-CoV-2 inoculation, co-cultivation of infected A549 together with PBMC gave rise to high levels of type I, II and III IFN, as well as increased amounts of TNF and IL-6. This is likely caused by direct cell-cell contact of pDCs and possibly other innate immune cells with infected A549, enabling sensing of and responding to infection by the immune cells as reported recently (61). In turn, this will sensitize virus sensing by the epithelial cells as we have shown above, leading to further increased IFN production. In fact, we found the majority of IFN-β and -λ to be dependent on RLR sensing in epithelial cells, highlighting the complex interplay of immune and epithelial cells in the establishing of a full-fledged antiviral response. It will be interesting to further dissect the contribution of epithelial vs. immune cells to the overall cytokine production in this setting in a future study. Of note, also here we found a significant tendency of stronger cytokine production when PBMC were from juvenile as compared to adult donors.

Our study has limitations and the proposed model clearly needs to be further tested in more authentic set-ups, ideally *in vivo*, in the future. In particular, our *in vitro* model is based on the lung adenocarcinoma-derived cell line A549, which might differ in various aspects from primary airway epithelial cells. Nonetheless, specifically the cell-intrinsic antiviral response is remarkably functional in A549 cells and they are widely accepted as a model system for this aspect. Furthermore, crude PBMC may be a poor surrogate for tissue resident or transiently patrolling immune cells found in the healthy mucosa. We tried to alleviate these limitations by cross-checking and validating our observations in scRNA-Seq data from nasal airway swabs. While our *in vitro* findings are compatible with accumulating human data from scRNA-Seq studies (19, 20), those *in vivo* data are only cross-sectional snapshots. Further, longitudinal experiments, e.g. in animal models such as the Syrian hamster (78, 79) or in mice (77), will be helpful in better understanding the complex interplay of immune and epithelial cells before and during infection.

Taken together, on the background of previous studies, our findings suggest a model of interplay of immune cells and epithelial cells in the context of SARS-CoV-2, and probably many other virus infections. Common pro-inflammatory cytokines such as TNF or IFN-γ likely constitutively present at low levels in the non-sterile environment of the airway mucosa critically tune the expression of virus sensing pathways in epithelial cells. In younger individuals up to adolescence, both an increased number of mucosal immune cells and their higher intrinsic propensity to produce such cytokines lead to an increased expression of MDA5 and RIG-I in epithelial cells as compared to adults. Under these conditions, SARS-CoV-2 infection is recognized more readily and leads to stronger induction of epithelial type I and III IFN and a more robust epithelial cell-intrinsic antiviral defense. The high number of immune cells and their elevated sensitivity in younger individuals in turn leads to a more ready detection of infected epithelial cells and consequently a stronger ensuing immune activation, amplifying the epithelial response and leading to a potent antiviral state of the tissue. This very likely affects the response to many if not all respiratory viruses. Seemingly at odds with our model, infections such as respiratory syncytial virus (RSV) or influenza virus are well-known to cause much higher morbidity amongst (young) children than in adults. However, it needs to be considered that those are widely circulating pathogens to which potent adaptive immune responses develop rapidly within the first few years of life (80, 81), providing powerful protection against severe disease throughout most of adulthood. For the pandemic SARS-CoV-2, in contrast, individuals of all age-classes were initially immuno-naïve. Interestingly, in the absence of adaptive immune responses, also RSV and influenza virus infections very likely cause more severe disease in adults as compared to children, as described for immunocompromised patients (82, 83) or in viral pandemics such as the 1918 “Spanish Flu”, during which large parts of the population initially were immuno-naïve (84). While this general age-trend, hence, is consistent with our proposed model, the clinically observed higher morbidity of RSV (and influenza) as compared to COVID-19 particularly in neonates and infants likely owes to a complex combination of virological, anatomical and immunological reasons.

In conclusion, our study sheds light on the vital contribution of immune cells to tissue homeostasis, in particular with regard to fine-tuning tissue sensitivity to virus infection. While this likely is of very broad and general relevance, we found it to be a hallmark difference between the airway mucosal innate immune response to SARS-CoV-2 infection in children and that in adults. A slight constitutive inflammatory milieu, mediated by immune cell-derived cytokines such as IFN-γ and TNF, leads to the IRF1-dependent upregulation of the crucial virus sensors RIG-I and MDA5, and thereby to a significantly increased overall sensitivity of the epithelial cell-intrinsic antiviral system. This enables the airway epithelium to more rapidly mount an efficacious and properly coordinated immune response, favoring successful containment of virus infection. Our results suggest it may be worthwhile exploring prophylactic strategies for SARS-CoV-2 and other respiratory infections aiming at mimicking the mucosal tissue homeostasis of school-age children, e.g. through inhalation of low-dose cytokine formulations (85).

## Acknowledgements

The authors want to thank Nadine Gillich for help in generating and characterizing the MDA5^KO^ and RIG-I/MDA5^DKO^ cell lines, Manina Günther for excellent help and guidance handling PBMC and *Ye*, and Dongsheng Yuan for help in scRNA-Seq data analysis. We are grateful to Ralf Bartenschlager for providing an outstanding research environment and critical discussions. We thank all patients and healthy volunteers who donated nasal swab samples or blood (PBMC) used in this study, and Sebastian Stricker for help with collection of nasal swabs. We further thank Maximilian Stich and Daniel Schnepf for critical reading of the manuscript. We apologize to all colleagues whose work we did not properly cite due to space limitations and/or our lack of oversight of the extensive amount of literature on COVID-19. Figures with schematic illustrations were made with biorender.com.

## Ethics Statement

Human material (PMBC) used in this study was obtained through studies approved by local ethics committees and conducted according to the declaration of Helsinki. Details (registration numbers) can be found in the methods section (Generation of PBMC).

## Author contributions

VGM, MD, SSB and SW performed experiments; VGM, SL, JL, SSB, IL and MB analyzed data and prepared visualizations; EMJ, KMD, RE, SA and AJ, JR and MAM provided essential material and/or indispensable intellectual input; VGM and MB conceived the study and wrote the manuscript; MB supervised and coordinated the study. All authors revised the article for important intellectual content and approved the final version.

---

**Supplementary Figure 1 (relates to Figure 1).**
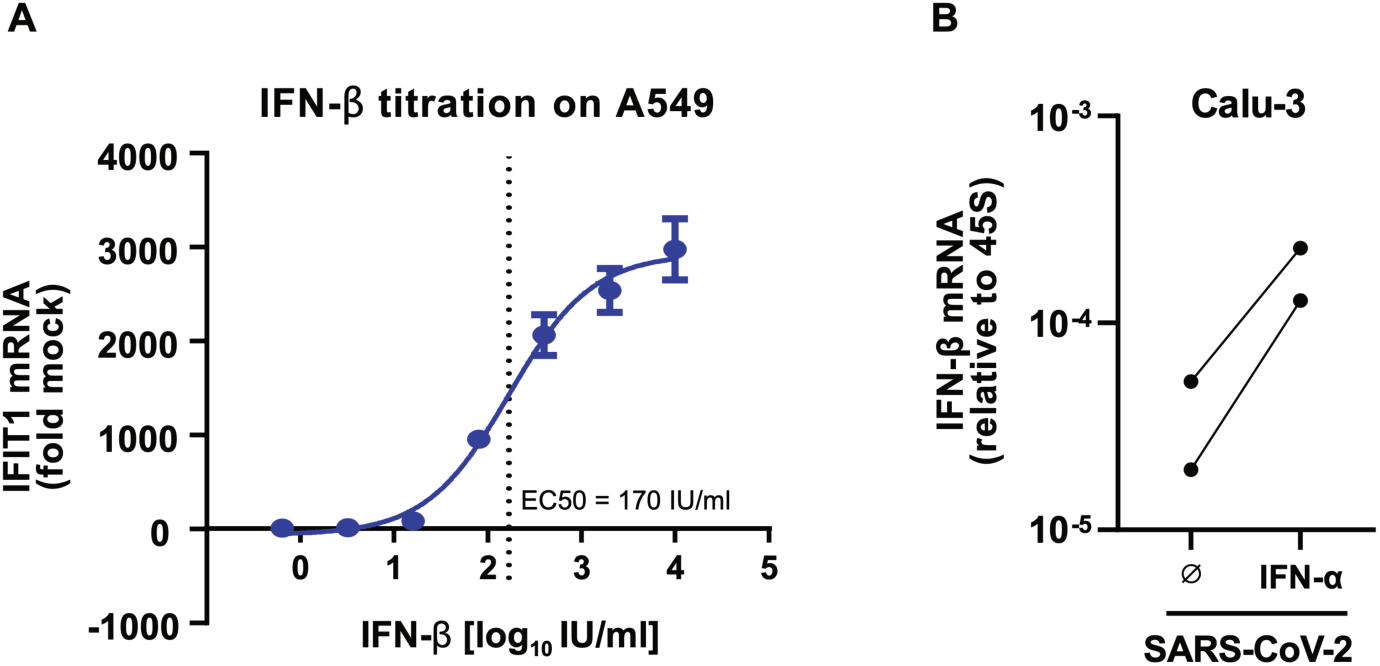
**(A)** A549 cells were mock-treated or primed with different dose of IFN-β for 16 hours. Cells were lysed and analyzed for IFIT1 gene expression by qRT-PCR. Shown are means ±SD of three technical replicates. **(B)** Calu-3 cells were mock-treated or primed with IFN-α (80 IU/ml) for 16 hours, followed by infection with SARS-CoV-2 (MOI 0.1). At 24 hours post infection, IFN-β expression was measured by qRT-PCR. Values were normalized to 45S rRNA levels and expressed relative to non-infected mock-treated cells. Quantification of two independent experiments are shown.

**Supplementary Figure 2 (relates to Figure 2).**
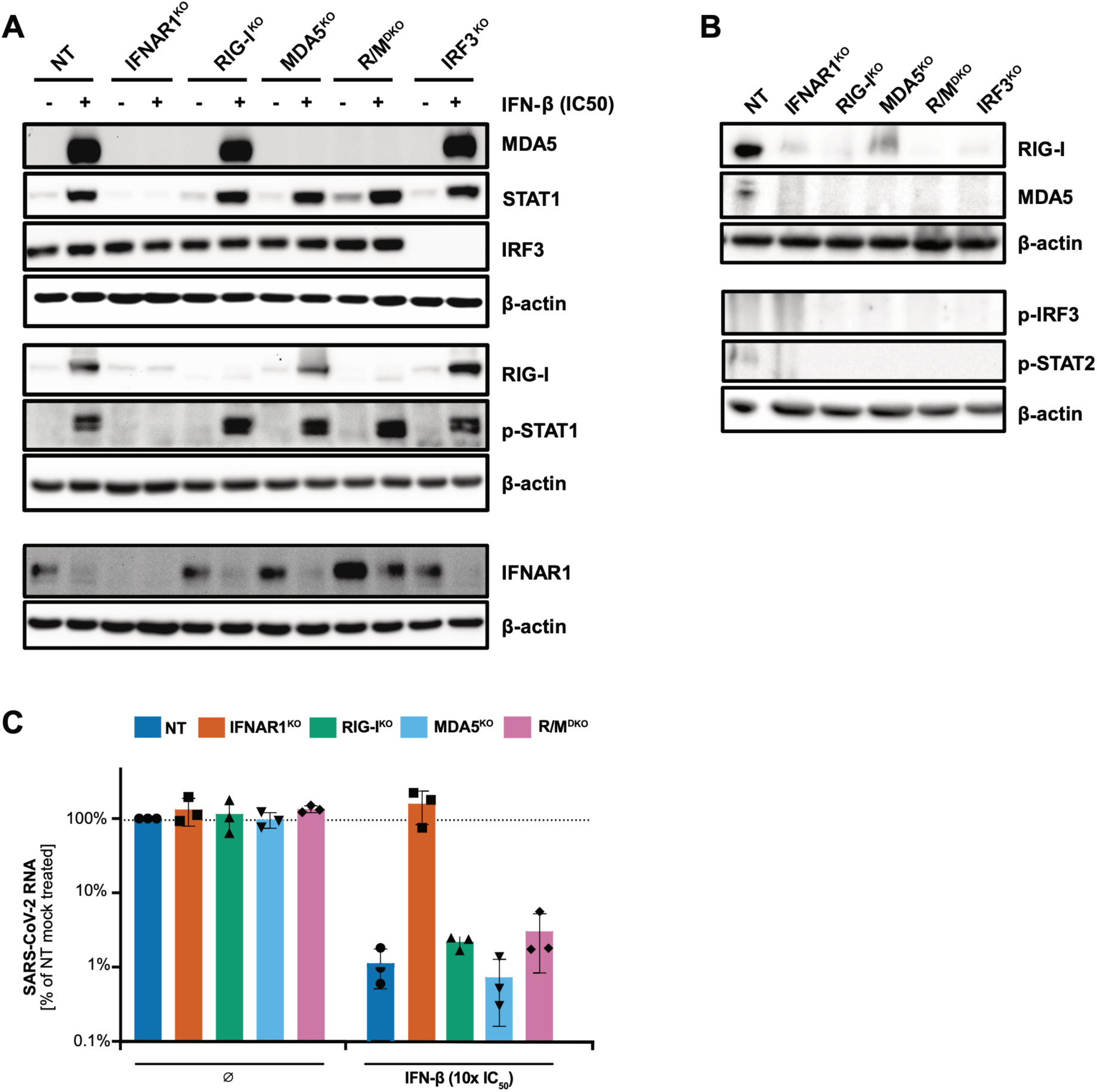
**(A)** A549^ACE2/TMPRSS2^ cells harboring the indicated CRISPR/Cas9-mediated functional KO were mock treated or primed with IFN-β (170 IU/ml) for 16 hours (NT: non-targeting as wildtype control; R/M^DKO^: RIG-I and MDA5 double KO). Cells were lysed and analyzed by immunoblotting using the indicated antibodies (“p” denotes phosphorylation specificity). Blot representative of two independent experiments. **(B)** Primed A549^ACE2/TMPRSS2^ (as in A, above) were mock-infected for 24 hours and lysed and analyzed by immunoblotting using the indicated antibodies. Blot representative of two independent experiments. (C) Mock-treated or IFN-β primed (10x IC50; 1,700 IU/ml) A549^ACE2/TMPRSS2^ cells harboring the indicated functional KOs (as in A) were infected with SARS-CoV-2 (MOI 0.1). At 24 hours post-infection, intracellular viral genome levels were determined by qRT-PCR using primers specific for ORF1. Values were normalized to 45S rRNA levels and expressed relative to non-primed wildtype (NT) cells.

**Supplementary Figure 3 (relates to Figure 4).**
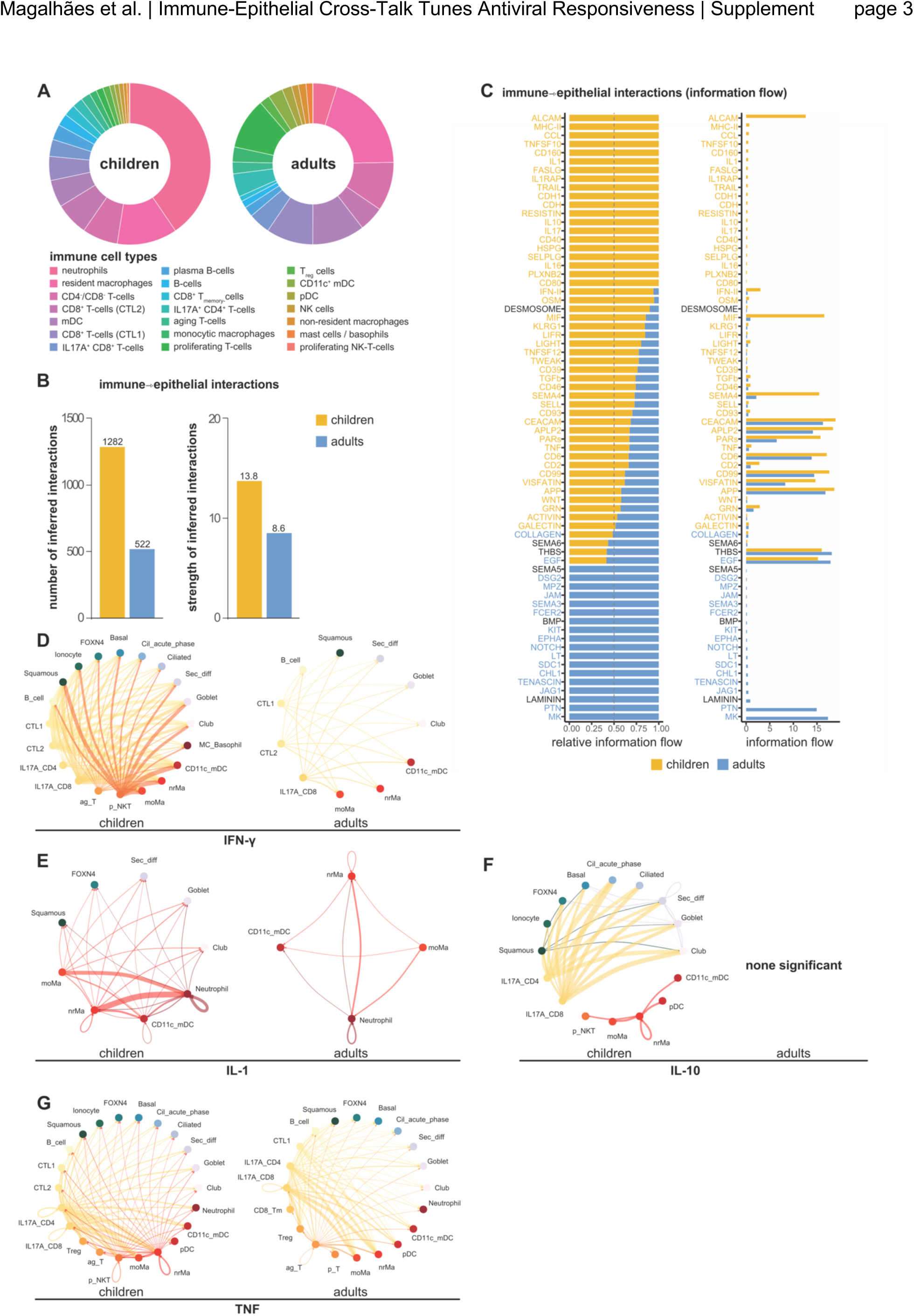
**(A)** Detailed composition of immune cell sub-populations in nasal swabs of healthy children and adults; total immune cell population set to 100% in each group separately. **(B)** Number (left) and strength (right) of ligand – receptor interactions in nasal swabs of children and adults (ligands expressed on immune cells, i.e. “senders”). **(C)** Immune – epithelial cell interactions mediated by the given ligand (on immune cells, i.e. “senders”). Relative (left) and absolute (right) information flow. **(D-G)** Interactions mediated by the specified cytokines expressed by immune cells (i.e. “senders”) mapped to individual cell types in children and adults. Interaction strength (information flow) coded by line thickness. **(B-G)** Two low-confidence cell types, “lowRNA” and “epithelial_lowRNA”, which showed little interaction with other cell types, were filtered and excluded for visual clarity.

**Supplementary Figure 4 (relates to Figure 5).**
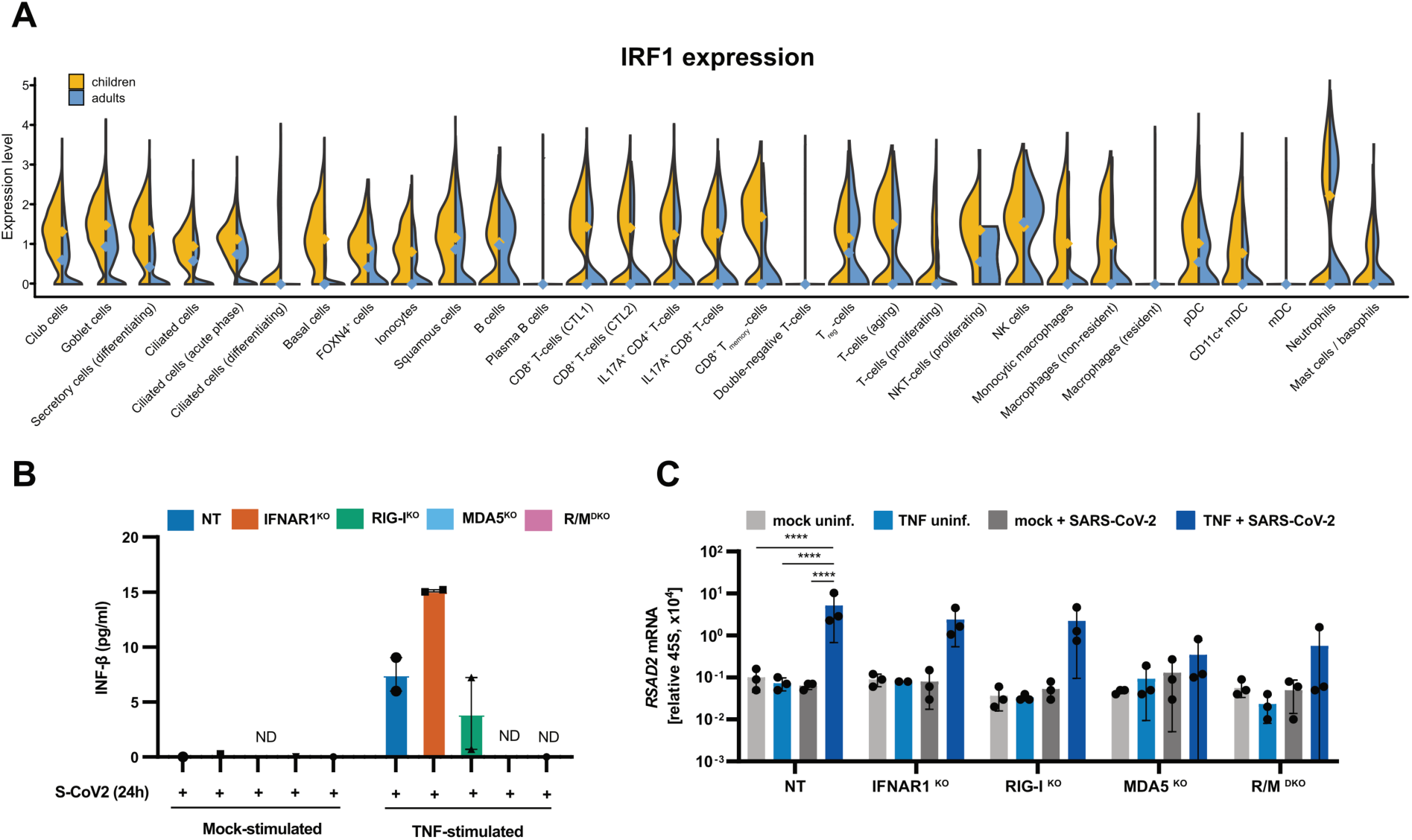
**(A)** Expression level of the transcription factor IRF1 in nasal swabs of children and adults across all identified cell types. Notch marks medians. **(B)** A549^ACE2/TMPRSS2^ cells harboring the indicated CRISPR/Cas9-mediated functional KO were mock treated or primed with TNF (10 ng/ml) for 8 hours followed by SARS-CoV-2 infection (MOI 1) (NT: non-targeting as wildtype control). At 24 hours post infection, culture supernatants were harvested and analyzed for secreted IFN-β. Data show the mean ±SD of two independent experiments. **(C)** A549^ACE2/TMPRSS2^ cells harboring the indicated CRISPR/Cas9-mediated functional KO were mock treated or primed with TNF (10 ng/ml) for 8 h followed by SARS-CoV-2 infection (MOI 1) (NT: non-targeting as wildtype control; R/M^DKO^: RIG-I and MDA5 double KO). At 24 h post infection, induction of IFN-β expression was determined by qRT-PCR. Values were normalized to 45S rRNA. Bars represent the mean ±SD of three independent experiments (individual experiments shown as dots). Statistical significance was tested by an unpaired one-tailed t-test. **** p < 0.0001.

**Supplementary Figure 5 (relates to Figure 6C).**
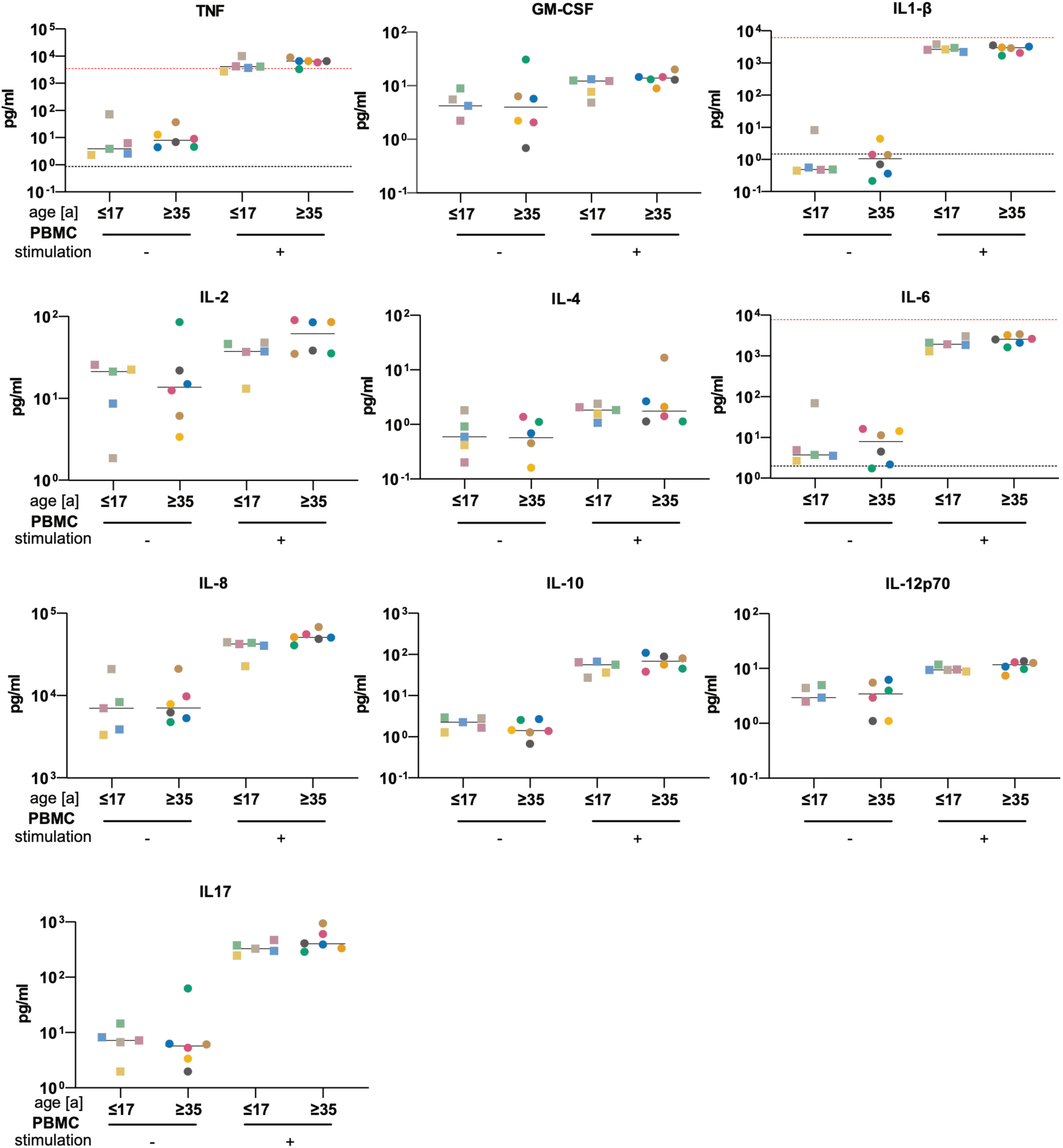
Peripheral blood mononuclear cells (PBMC) isolated from healthy juvenile (14 – 17 years) and adult (35 – 60 years) donors were mock treated or exposed to live Gram-negative bacteria (*Yersinia enterocolitica*) for 1 hour, after which bacteria were killed by antibiotic (gentamicin) treatment. Upon further incubation for 24 hours, culture supernatants were harvested and analyzed. Values for each donor are shown individually (5 juvenile, 6 adult donors), color identifies donor across all measurements. Median shown as solid black line. Dashed line in black indicates lower limit of detection, whereas dashed line in red denotes the upper limit of detection of the indicated analyte.

**Supplementary Figure 6 (relates to Figure 6 E).**
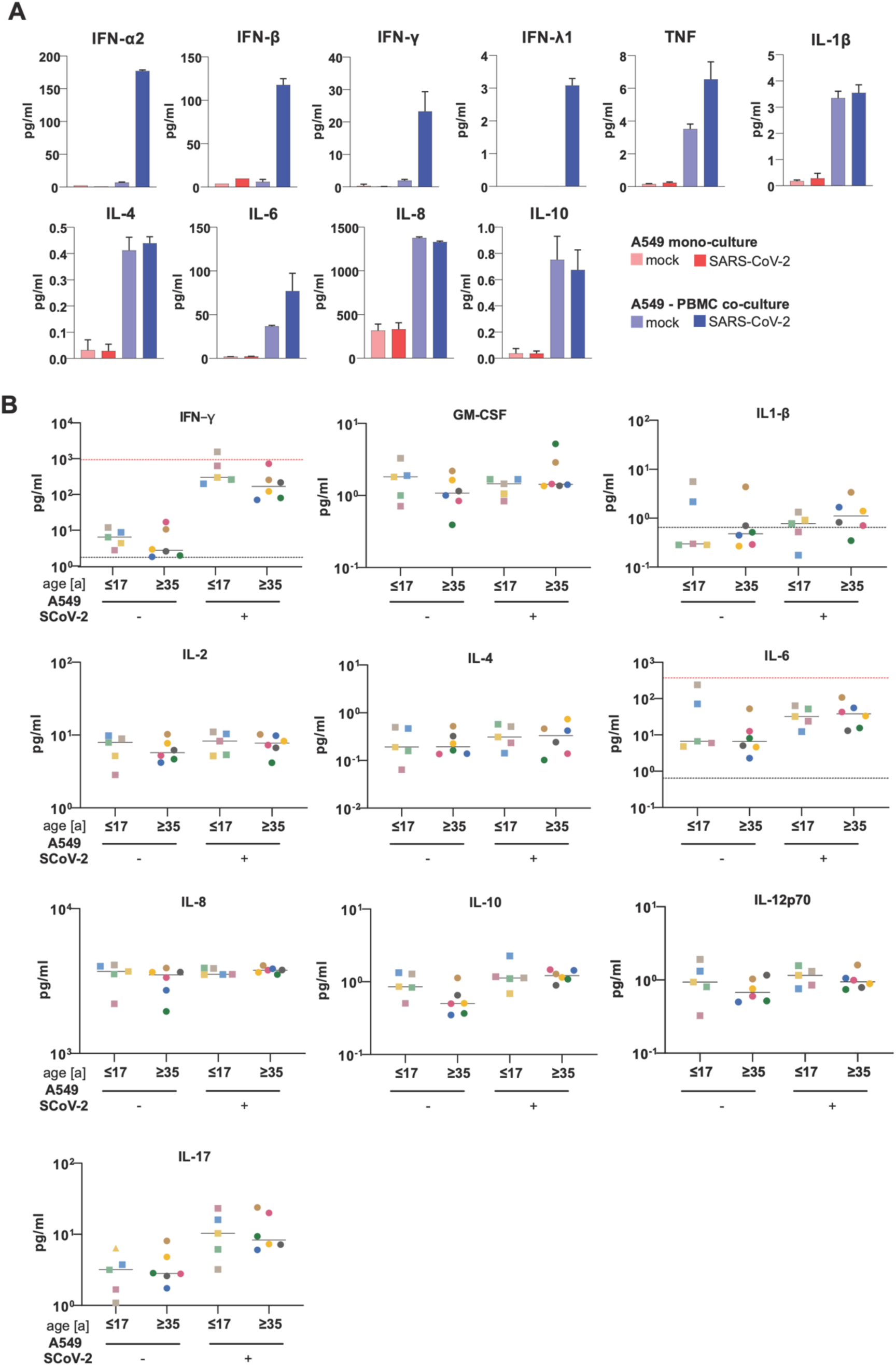
**(A)** A549 cells were infected with SARS-CoV-2 (MOI 0.1) or mock-infected for 1 hour before adding PBMC to the culture. After 24 h of co-culturing, supernatants were harvested and subjected to cytokine multiplex measurement. Data show the mean ±SD of three technical replicates. **(B)** A549 cells were infected with SARS-CoV-2 (MOI 0.1) for 8 hours before adding isolated PBMC from healthy juvenile (14-17 years) and adult (35-60 years) donors. After 24 hours of co-culturing, supernatants were harvested and subjected to cytokine multiplex measurement. Values for each donor are shown individually (5 juvenile, 6 adult donors), color identifies donor across all measurements. Median shown as solid black line. Dashed line in black indicates lower limit of detection, whereas dashed line in red denotes the upper limit of detection of the indicated analyte.

**Supplementary Figure 7 (relates to Figure 6).**
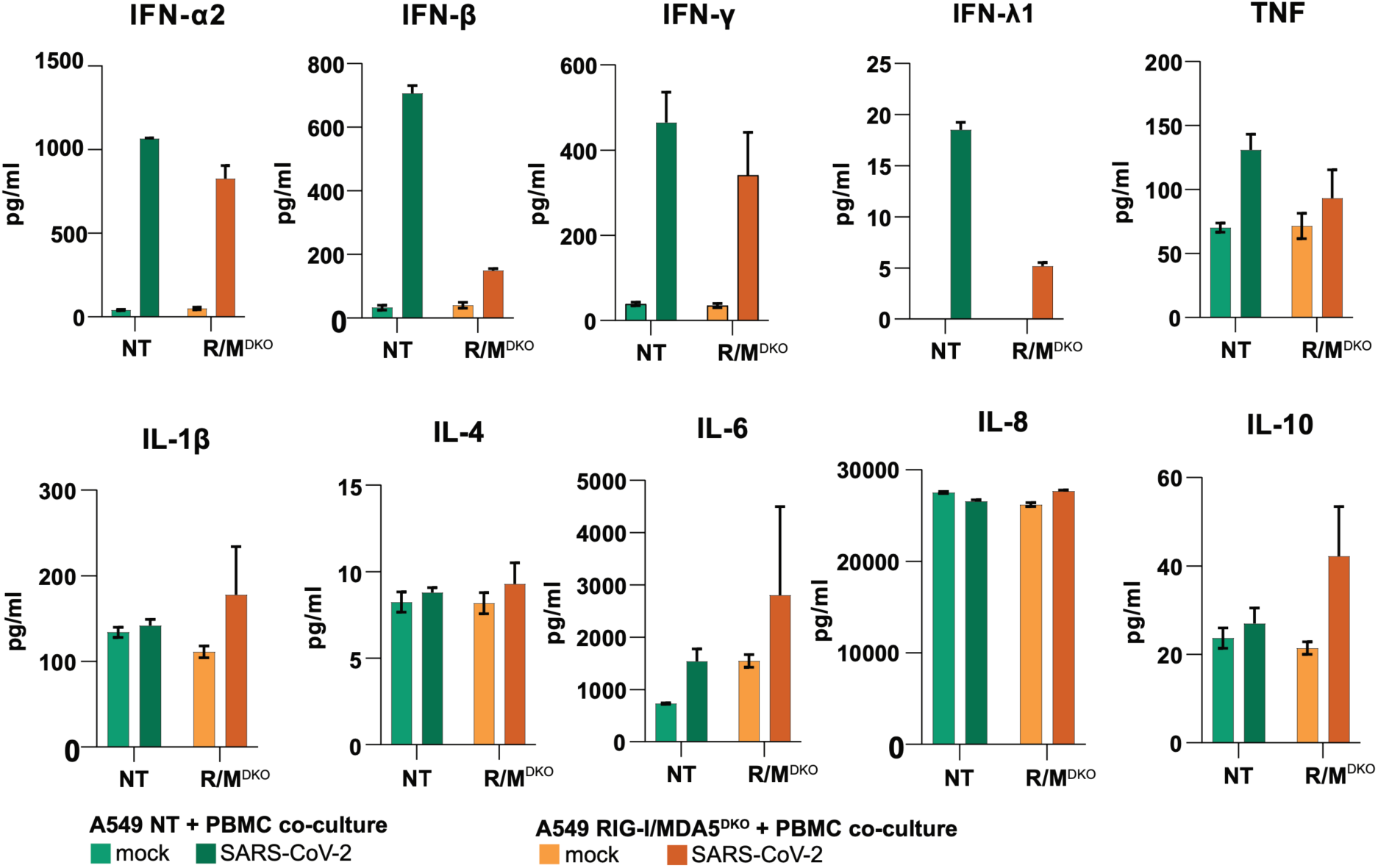
A549 NT (as wildtype controls) or A549 RIG-I/MDA5^DKO^ (R/M^DKO^) cells were infected with SARS-CoV-2 (MOI 0.1) for 1 hour before adding PBMC to the culture. After 24 hours of co-culturing, supernatants were harvested and subjected to cytokine multiplex measurement. Data show the mean ±SD of three technical replicates.

**Supplementary Figure 8 (relates to materials and methods).**
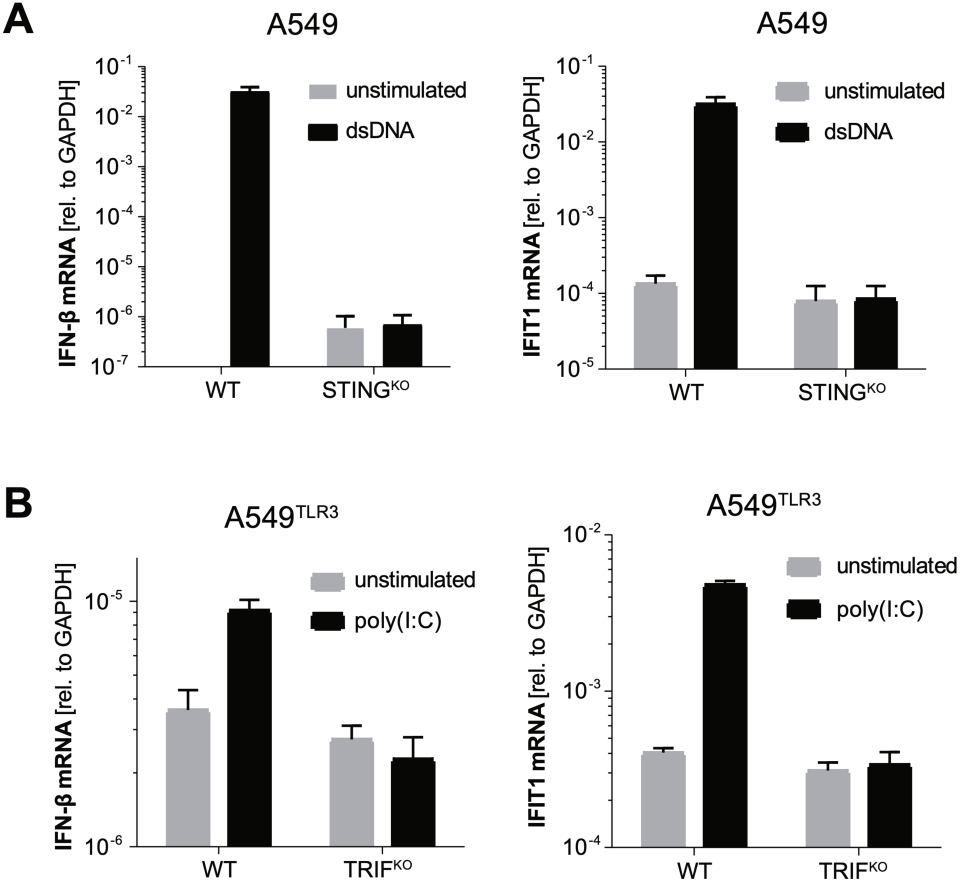
Transcription levels of IFN-β1 and IFIT1 in **(A)** STING^KO^ and **(B)** TRIF^KO^ clones were assessed by qRT-PCR after specific pathway stimulation. For the cGAS/STING pathway dsDNA (herring sperm DNA) was transfected. For the TLR3/TRIF pathway, poly(I:C) was added to the culture medium; due to very low baseline TLR3 expression in A549, cells were stably transduced with TLR3. Data show the mean ±SD of three technical replicates.

## References

1. Guan WJ, Ni ZY, Hu Y, Liang WH, Ou CQ, He JX, et al. Clinical Characteristics of Coronavirus Disease 2019 in China. N Engl J Med. 2020;382(18):1708–20.

2. Chen N, Zhou M, Dong X, Qu J, Gong F, Han Y, et al. Epidemiological and clinical characteristics of 99 cases of 2019 novel coronavirus pneumonia in Wuhan, China: a descriptive study. The Lancet. 2020;395(10223):507–13.

3. Zhou F, Yu T, Du R, Fan G, Liu Y, Liu Z, et al. Clinical course and risk factors for mortality of adult inpatients with COVID-19 in Wuhan, China: a retrospective cohort study. The Lancet. 2020;395(10229):1054–62.

4. Wang H, Paulson KR, Pease SA, Watson S, Comfort H, Zheng P, et al. Estimating excess mortality due to the COVID-19 pandemic: a systematic analysis of COVID-19-related mortality, 2020–21. The Lancet. 2022;399(10334):1513–36.

5. Ng WH, Tipih T, Makoah NA, Vermeulen JG, Goedhals D, Sempa JB, et al. Comorbidities in SARS-CoV-2 Patients: a Systematic Review and Meta-Analysis. mBio. 2021;12(1).

6. O’Driscoll M, Ribeiro Dos Santos G, Wang L, Cummings DAT, Azman AS, Paireau J, et al. Age-specific mortality and immunity patterns of SARS-CoV-2. Nature. 2021;590(7844):140–5.

7. Jones TC, Biele G, Mühlemann B, Veith T, Schneider J, Beheim-Schwarzbach J, et al. Estimating infectiousness throughout SARS-CoV-2 infection course. Science. 2021;373(6551).

8. Toh ZQ, Anderson J, Mazarakis N, Neeland M, Higgins RA, Rautenbacher K, et al. Comparison of Seroconversion in Children and Adults With Mild COVID-19. JAMA Netw Open. 2022;5(3):e221313.

9. Zhang Q, Bastard P, Liu Z, Le Pen J, Moncada-Velez M, Chen J, et al. Inborn errors of type I IFN immunity in patients with life-threatening COVID-19. Science. 2020;370(6515).

10. Bastard P, Rosen LB, Zhang Q, Michailidis E, Hoffmann HH, Zhang Y, et al. Autoantibodies against type I IFNs in patients with life-threatening COVID-19. Science. 2020;370(6515).

11. Lopez J, Mommert M, Mouton W, Pizzorno A, Brengel-Pesce K, Mezidi M, et al. Early nasal type I IFN immunity against SARS-CoV-2 is compromised in patients with autoantibodies against type I IFNs. J Exp Med. 2021;218(10).

12. Lee JS, Park S, Jeong HW, Ahn JY, Choi SJ, Lee H, et al. Immunophenotyping of COVID-19 and influenza highlights the role of type I interferons in development of severe COVID-19. Science immunology. 2020;5(49).

13. Lucas C, Wong P, Klein J, Castro TBR, Silva J, Sundaram M, et al. Longitudinal analyses reveal immunological misfiring in severe COVID-19. Nature. 2020;584(7821):463–9.

14. Park A, Iwasaki A. Type I and Type III Interferons - Induction, Signaling, Evasion, and Application to Combat COVID-19. Cell host & microbe. 2020;27(6):870–8.

15. Wong LR, Perlman S. Immune dysregulation and immunopathology induced by SARS-CoV-2 and related coronaviruses - are we our own worst enemy? Nat Rev Immunol. 2022;22(1):47–56.

16. Sposito B, Broggi A, Pandolfi L, Crotta S, Clementi N, Ferrarese R, et al. The interferon landscape along the respiratory tract impacts the severity of COVID-19. Cell. 2021;184(19):4953–68.e16.

17. Cheemarla NR, Watkins TA, Mihaylova VT, Wang B, Zhao D, Wang G, et al. Dynamic innate immune response determines susceptibility to SARS-CoV-2 infection and early replication kinetics. J Exp Med. 2021;218(8).

18. Blanco-Melo D, Nilsson-Payant BE, Liu WC, Uhl S, Hoagland D, Moller R, et al. Imbalanced Host Response to SARS-CoV-2 Drives Development of COVID-19. Cell. 2020;181(5):1036–45 e9.

19. Yoshida M, Worlock KB, Huang N, Lindeboom RGH, Butler CR, Kumasaka N, et al. Local and systemic responses to SARS-CoV-2 infection in children and adults. Nature. 2022;602(7896):321–7.

20. Loske J, Rohmel J, Lukassen S, Stricker S, Magalhaes VG, Liebig J, et al. Pre-activated antiviral innate immunity in the upper airways controls early SARS-CoV-2 infection in children. Nat Biotechnol. 2022;40(3):319–24.

21. Lamers MM, Haagmans BL. SARS-CoV-2 pathogenesis. Nat Rev Microbiol. 2022;20(5):270–84.

22. Chu H, Chan JF-W, Yuen TT-T, Shuai H, Yuan S, Wang Y, et al. Comparative tropism, replication kinetics, and cell damage profiling of SARS-CoV-2 and SARS-CoV with implications for clinical manifestations, transmissibility, and laboratory studies of COVID-19: an observational study. The Lancet Microbe. 2020;1(1):e14–e23.

23. Neufeldt CJ, Cerikan B, Cortese M, Frankish J, Lee JY, Plociennikowska A, et al. SARS-CoV-2 infection induces a pro-inflammatory cytokine response through cGAS-STING and NF-kappaB. Commun Biol. 2022;5(1):45.

24. Hayn M, Hirschenberger M, Koepke L, Nchioua R, Straub JH, Klute S, et al. Systematic functional analysis of SARS-CoV-2 proteins uncovers viral innate immune antagonists and remaining vulnerabilities. Cell Rep. 2021;35(7):109126.

25. Lee JH, Koepke L, Kirchhoff F, Sparrer KMJ. Interferon antagonists encoded by SARS-CoV-2 at a glance. Medical microbiology and immunology. 2022:1–7.

26. Yin X, Riva L, Pu Y, Martin-Sancho L, Kanamune J, Yamamoto Y, et al. MDA5 Governs the Innate Immune Response to SARS-CoV-2 in Lung Epithelial Cells. Cell Rep. 2021;34(2):108628.

27. Sampaio NG, Chauveau L, Hertzog J, Bridgeman A, Fowler G, Moonen JP, et al. The RNA sensor MDA5 detects SARS-CoV-2 infection. Scientific Reports. 2021;11(1):13638.

28. Rebendenne A, Valadão ALC, Tauziet M, Maarifi G, Bonaventure B, McKellar J, et al. SARS-CoV-2 triggers an MDA-5-dependent interferon response which is unable to control replication in lung epithelial cells. J Virol. 2021;95(8).

29. Nilsson-Payant BE, Uhl S, Grimont A, Doane AS, Cohen P, Patel RS, et al. The NF-kappaB Transcriptional Footprint Is Essential for SARS-CoV-2 Replication. J Virol. 2021;95(23):e0125721.

30. Krischuns T, Günl F, Henschel L, Binder M, Willemsen J, Schloer S, et al. Phosphorylation of TRI M28 Enhances the Expression of IFN-β and Proinflammatory Cytokines During HPAIV Infection of Human Lung Epithelial Cells. 2018;9.

31. Willemsen J, Wicht O, Wolanski JC, Baur N, Bastian S, Haas DA, et al. Phosphorylation-Dependent Feedback Inhibition of RIG-I by DAPK1 Identified by Kinome-wide siRNA Screening. Molecular cell. 2017;65(3):403–15.e8.

32. Plociennikowska A, Frankish J, Moraes T, Del Prete D, Kahnt F, Acuna C, et al. TLR3 activation by Zika virus stimulates inflammatory cytokine production which dampens the antiviral response induced by RIG-I-like receptors. J Virol. 2021.

33. Urban C, Welsch H, Heine K, Wüst S, Haas DA, Dächert C, et al. Persistent Innate Immune Stimulation Results in IRF3-Mediated but Caspase-Independent Cytostasis. Viruses. 2020;12(6).

34. Wüst S, Schad P, Burkart S, Binder M. Comparative Analysis of Six IRF Family Members in Alveolar Epithelial Cell-Intrinsic Antiviral Responses. 2021;10(10):2600.

35. Sanjana NE, Shalem O, Zhang F. Improved vectors and genome-wide libraries for CRISPR screening. Nature methods. 2014;11(8):783–4.

36. Haddad A, Janda A, Renk H, Stich M, Frieh P, Kaier K, et al. Long COVID symptoms in exposed and infected children, adolescents and their parents one year after SARS-CoV-2 infection: A prospective observational cohort study. EBioMedicine. 2022;84:104245.

37. Renk H, Dulovic A, Seidel A, Becker M, Fabricius D, Zernickel M, et al. Robust and durable serological response following pediatric SARS-CoV-2 infection. Nat Commun. 2022;13(1):128.

38. Stich M, Elling R, Renk H, Janda A, Garbade SF, Müller B, et al. Transmission of Severe Acute Respiratory Syndrome Coronavirus 2 in Households with Children, Southwest Germany, May-August 2020. Emerging infectious diseases. 2021;27(12):3009–19.

39. Burkart SS, Schweinoch D, Frankish J, Sparn C, Wüst S, Urban C, et al. High Resolution Kinetic Characterization and Dynamic Mathematical Modeling of the RIG-I Signaling Pathway and the Antiviral Responses. 2022:2022.08.05.502818.

40. Binder M, Eberle F, Seitz S, Mücke N, Hüber CM, Kiani N, et al. Molecular Mechanism of Signal Perception and Integration by the Innate Immune Sensor Retinoic Acid-inducible Gene-I (RIG-I)*. Journal of Biological Chemistry. 2011;286(31):27278–87.

41. Jin S, Guerrero-Juarez CF, Zhang L, Chang I, Ramos R, Kuan CH, et al. Inference and analysis of cell-cell communication using CellChat. Nat Commun. 2021;12(1):1088.

42. Shilts J, Severin Y, Galaway F, Müller-Sienerth N, Chong Z-S, Pritchard S, et al. A physical wiring diagram for the human immune system. Nature. 2022;608(7922):397–404.

43. Benjamini Y, Hochberg Y. Controlling the False Discovery Rate: A Practical and Powerful Approach to Multiple Testing. Journal of the Royal Statistical Society Series B (Methodological). 1995;57(1):289–300.

44. Chu H, Chan JF, Wang Y, Yuen TT, Chai Y, Hou Y, et al. Comparative Replication and Immune Activation Profiles of SARS-CoV-2 and SARS-CoV in Human Lungs: An Ex Vivo Study With Implications for the Pathogenesis of COVID-19. Clinical infectious diseases : an official publication of the Infectious Diseases Society of America. 2020;71(6):1400–9.

45. Neeland MR, Bannister S, Clifford V, Nguyen J, Dohle K, Overmars I, et al. Children and Adults in a Household Cohort Study Have Robust Longitudinal Immune Responses Following SARS-CoV-2 Infection or Exposure. Front Immunol. 2021;12:741639.

46. Vono M, Huttner A, Lemeille S, Martinez-Murillo P, Meyer B, Baggio S, et al. Robust innate responses to SARS-CoV-2 in children resolve faster than in adults without compromising adaptive immunity. Cell Rep. 2021;37(1):109773.

47. Thorne LG, Bouhaddou M, Reuschl AK, Zuliani-Alvarez L, Polacco B, Pelin A, et al. Evolution of enhanced innate immune evasion by SARS-CoV-2. Nature. 2022;602(7897):487–95.

48. Thorne LG, Reuschl AK, Zuliani-Alvarez L, Whelan MVX, Turner J, Noursadeghi M, et al. SARS-CoV-2 sensing by RIG-I and MDA5 links epithelial infection to macrophage inflammation. EMBO J. 2021;40(15):e107826.

49. Khanmohammadi S, Rezaei N. Role of Toll-like receptors in the pathogenesis of COVID-19. Journal of medical virology. 2021;93(5):2735–9.

50. Asano T, Boisson B, Onodi F, Matuozzo D, Moncada-Velez M, Maglorius Renkilaraj MRL, et al. X-linked recessive TL R7 deficiency in ∼1% of men under 60 years old with life-threatening COVID-19. 2021;6(62):eabl4348.

51. Sposito B, Broggi A, Pandolfi L, Crotta S, Clementi N, Ferrarese R, et al. The interferon landscape along the respiratory tract impacts the severity of COVID-19. Cell. 2021;184(19):4953–68 e16.

52. Larsen SB, Cowley CJ, Fuchs E. Epithelial cells: liaisons of immunity. Curr Opin Immunol. 2020;62:45–53.

53. Hewitt RJ, Lloyd CM. Regulation of immune responses by the airway epithelial cell landscape. Nature Reviews Immunology. 2021;21(6):347–62.

54. Willemsen J, Neuhoff MT, Hoyler T, Noir E, Tessier C, Sarret S, et al. TNF leads to mtDNA release and cGAS/STING-dependent interferon responses that support inflammatory arthritis. Cell Rep. 2021;37(6):109977.

55. Aarreberg LD, Esser-Nobis K, Driscoll C, Shuvarikov A, Roby JA, Gale M, Jr. Interleukin-1β Induces mtDNA Release to Activate Innate Immune Signaling via cGAS-STING. Molecular cell. 2019;74(4):801–15.e6.

56. Panda D, Gjinaj E, Bachu M, Squire E, Novatt H, Ozato K, et al. IRF1 Maintains Optimal Constitutive Expression of Antiviral Genes and Regulates the Early Antiviral Response. Front Immunol. 2019;10:1019.

57. Yarilina A, Park-Min KH, Antoniv T, Hu X, Ivashkiv LB. TNF activates an IRF1-dependent autocrine loop leading to sustained expression of chemokines and STAT1-dependent type I interferon-response genes. Nat Immunol. 2008;9(4):378–87.

58. Forero A, Ozarkar S, Li H, Lee CH, Hemann EA, Nadjsombati MS, et al. Differential Activation of the Transcription Factor IRF1 Underlies the Distinct Immune Responses Elicited by Type I and Type III Interferons. Immunity. 2019;51(3):451–64.e6.

59. Lee SJ, Jang BC, Lee SW, Yang YI, Suh SI, Park YM, et al. Interferon regulatory factor-1 is prerequisite to the constitutive expression and IFN-gamma-induced upregulation of B7-H1 (CD274). FEBS letters. 2006;580(3):755–62.

60. Murtas D, Maric D, De Giorgi V, Reinboth J, Worschech A, Fetsch P, et al. IRF-1 responsiveness to IFN-γ predicts different cancer immune phenotypes. British journal of cancer. 2013;109(1):76–82.

61. Venet M, Sa Ribeiro M, Décembre E, Bellomo A, Joshi G, Villard M, et al. Severe COVID-19 patients have impaired plasmacytoid dendritic cell-mediated control of SARS-CoV-2-infected cells. 2021:2021.09.01.21262969.

62. Newton AH, Cardani A, Braciale TJ. The host immune response in respiratory virus infection: balancing virus clearance and immunopathology. Semin Immunopathol. 2016;38(4):471–82.

63. Rai KR, Shrestha P, Yang B, Chen Y, Liu S, Maarouf M, et al. Acute Infection of Viral Pathogens and Their Innate Immune Escape. Front Microbiol. 2021;12:672026.

64. Desai N, Neyaz A, Szabolcs A, Shih AR, Chen JH, Thapar V, et al. Temporal and spatial heterogeneity of host response to SARS-CoV-2 pulmonary infection. Nat Commun. 2020;11(1):6319.

65. Katsura H, Sontake V, Tata A, Kobayashi Y, Edwards CE, Heaton BE, et al. Human Lung Stem Cell-Based Alveolospheres Provide Insights into SARS-CoV-2-Mediated Interferon Responses and Pneumocyte Dysfunction. Cell Stem Cell. 2020;27(6):890–904 e8.

66. Stanifer ML, Kee C, Cortese M, Zumaran CM, Triana S, Mukenhirn M, et al. Critical Role of Type III Interferon in Controlling SARS-CoV-2 Infection in Human Intestinal Epithelial Cells. Cell Rep. 2020;32(1):107863.

67. Lokugamage KG, Hage A, de Vries M, Valero-Jimenez AM, Schindewolf C, Dittmann M, et al. Type I interferon susceptibility distinguishes SARS-CoV-2 from SARS-CoV. bioRxiv. 2020.

68. Vanderheiden A, Ralfs P, Chirkova T, Upadhyay AA, Zimmerman MG, Bedoya S, et al. Type I and Type III Interferons Restrict SARS-CoV-2 Infection of Human Airway Epithelial Cultures. J Virol. 2020;94(19).

69. Felgenhauer U, Schoen A, Gad HH, Hartmann R, Schaubmar AR, Failing K, et al. Inhibition of SARS-CoV-2 by type I and type III interferons. The Journal of biological chemistry. 2020;295(41):13958–64.

70. Meisel C, Akbil B, Meyer T, Lankes E, Corman VM, Staudacher O, et al. Mild COVID-19 despite autoantibodies against type I IFNs in autoimmune polyendocrine syndrome type 1. J Clin Invest. 2021;131(14).

71. Yamada T, Sato S, Sotoyama Y, Orba Y, Sawa H, Yamauchi H, et al. RIG-I triggers a signaling-abortive anti-SARS-CoV-2 defense in human lung cells. Nat Immunol. 2021;22(7):820–8.

72. Molony RD, Nguyen JT, Kong Y, Montgomery RR, Shaw AC, Iwasaki A. Aging impairs both primary and secondary RIG-I signaling for interferon induction in human monocytes. Sci Signal. 2017;10(509).

73. Wu X, Dao Thi VL, Huang Y, Billerbeck E, Saha D, Hoffmann HH, et al. Intrinsic Immunity Shapes Viral Resistance of Stem Cells. Cell. 2018;172(3):423–38 e25.

74. Crow YJ, Stetson DB. The type I interferonopathies: 10 years on. Nat Rev Immunol. 2022;22(8):471–83.

75. Venkatesh D, Ernandez T, Rosetti F, Batal I, Cullere X, Luscinskas FW, et al. Endothelial TNF receptor 2 induces IRF1 transcription factor-dependent interferon-beta autocrine signaling to promote monocyte recruitment. Immunity. 2013;38(5):1025–37.

76. Shaw AC, Goldstein DR, Montgomery RR. Age-dependent dysregulation of innate immunity. Nature Reviews Immunology. 2013;13(12):875–87.

77. Beer J, Crotta S, Breithaupt A, Ohnemus A, Becker J, Sachs B, et al. Impaired immune response drives age-dependent severity of COVID-19. J Exp Med. 2022;219(12).

78. Nouailles G, Wyler E, Pennitz P, Postmus D, Vladimirova D, Kazmierski J, et al. Temporal omics analysis in Syrian hamsters unravel cellular effector responses to moderate COVID-19. Nature Communications. 2021;12(1):4869.

79. Francis ME, Goncin U, Kroeker A, Swan C, Ralph R, Lu Y, et al. SARS-CoV-2 infection in the Syrian hamster model causes inflammation as well as type I interferon dysregulation in both respiratory and non-respiratory tissues including the heart and kidney. PLoS Pathog. 2021;17(7):e1009705.

80. Bodewes R, de Mutsert G, van der Klis FR, Ventresca M, Wilks S, Smith DJ, et al. Prevalence of antibodies against seasonal influenza A and B viruses in children in Netherlands. Clin Vaccine Immunol. 2011;18(3):469–76.

81. Andeweg SP, Schepp RM, van de Kassteele J, Mollema L, Berbers GAM, van Boven M. Population-based serology reveals risk factors for RSV infection in children younger than 5 years. Sci Rep. 2021;11(1):8953.

82. Chatzis O, Darbre S, Pasquier J, Meylan P, Manuel O, Aubert JD, et al. Burden of severe RSV disease among immunocompromised children and adults: a 10 year retrospective study. BMC Infect Dis. 2018;18(1):111.

83. Whitley RJ, Monto AS. Prevention and treatment of influenza in high-risk groups: children, pregnant women, immunocompromised hosts, and nursing home residents. J Infect Dis. 2006;194 Suppl 2:S133–8.

84. Ahmed R, Oldstone MB, Palese P. Protective immunity and susceptibility to infectious diseases: lessons from the 1918 influenza pandemic. Nat Immunol. 2007;8(11):1188–93.

85. Meng Z, Wang T, Chen L, Chen X, Li L, Qin X, et al. The Effect of Recombinant Human Interferon Alpha Nasal Drops to Prevent COVID-19 Pneumonia for Medical Staff in an Epidemic Area. Current topics in medicinal chemistry. 2021;21(10):920–7.

